# Nucleus Accumbens Local Circuit for Cue-Dependent Aversive Learning

**DOI:** 10.1101/2023.02.06.527338

**Authors:** Andrew Belilos, Cortez Gray, Christie Sanders, Destiny Black, Elizabeth Mays, Christopher Richie, Ayesha Sengupta, Holly Hake, T. Chase Francis

## Abstract

Response to threatening environmental stimuli requires detection and encoding of important environmental features that dictate threat. Aversive events are highly salient which promotes associative learning about stimuli that signal this threat. The nucleus accumbens is uniquely positioned to process this salient, aversive information and promote motivated output, through plasticity on the major projection neurons in the brain area. We uncovered a nucleus accumbens core local circuit whereby excitatory plasticity facilitates learning and recall of discrete aversive cues. We demonstrate that putative nucleus accumbens substance P release and long-term excitatory plasticity on dopamine 2 receptor expressing projection neurons is required for cue-dependent fear learning. Additionally, we found fear learning and recall were dependent on distinct projection-neuron subtypes. Our work demonstrates a critical role for Nucleus Accumbens substance P in cue-dependent aversive learning.

## Introduction

Aversive stimuli are inherently salient, as animals need to recognize these threats to survive. Consequently, aversive stimuli elicit rapid learning about and response to cues that predict aversive outcomes. Canonically, the major projection neurons of the Nucleus Accumbens (NAc) are studied in terms of motivational or incentive salience, a cognitive process that facilitates approach or avoidance in response to a stimulus. These responses, however, require recognition of a salient stimulus to promote these motivated outcomes. The NAc responds to salient cues in humans ^1^, is required for aversive learning ^2,3^, which is cue-dependent in the NAc core ^3–5^, suggesting the NAc core may facilitate Pavlovian learning independently of the motivated response. If true, it is likely consolidation and subsequent recall of these salient events requires long-term plasticity to facilitate persistent memories. These NAc-dependent synaptic mechanisms to enable recognition of threatening, aversive stimuli have yet to be determined.

Within the NAc, medium spiny projection neuron (MSN) subtypes are thought to promote opposing outcomes, with dopamine 1 (D1) receptor MSN activity driving rewarding and positive outcomes, and dopamine 2 (D2) receptor MSN activity driving aversive outcomes, suppressed reward, and punishment ^6–15^. This understanding has been updated and it is now known MSN subtype activity participates in more complex and pivotal roles in guiding learned behavior. In addition, NAc core D2-MSNs have been shown to be required for discrimination ^16^ and reversal learning ^17,18^, and reward ^19–21^. These additional known roles of D2-MSNs may hinge on the ability of D2-MSNs to alter their response to salient stimuli. Thereby, stimulus-dependent activity (e.g., through potentiation of excitatory transmission on D2-MSNs), during learning, may provide a means of facilitating learning about aversion associated stimuli.

We recently uncovered a mechanism in which glutamatergic long-term excitatory potentiation (LTP) is elicited on D2-MSNs exclusively by activity of D1-MSNs ^22^ and required the activation of cholinergic interneurons by release of the neuropeptide substance P. Both acetylcholine and substance P are released by salient stimuli including drugs of abuse and aversive stimuli ^23,24^ and these neuromodulators contribute to many of the same behaviors as activation of D2-MSNs. Additionally, phasic activity changes of cholinergic neurons in response to cue-dependent associative outcomes have been observed in primates ^25,26^ and rodents ^27^. Like D2-MSN neuronal activity, striatal acetylcholine is necessary for behavioral flexibility ^28^, discrimination ^29^, reversal learning ^30^, and aids in the expression of cue-motivated behavior ^31^. These similarities, and the fact that this circuit promotes plasticity position this circuit as a likely candidate for signaling aversion-related outcomes. Therefore, we hypothesized this NAc core substance P is necessary for supporting aversive fear learning through plasticity on D2-MSNs.

In this paper, we aimed to determine synaptic and circuit mechanisms which underpin NAc core dependent aversive learning. Using pharmacology and gene knockout technology, we found substance P receptors in cholinergic neurons were required for cue-dependent aversive learning, in a fear conditioning context. These findings were corroborated through enhanced D1-MSN activity and acetylcholine release in response to aversive foot shocks and D2-MSN activity in response to learned cues. Disrupting this circuit at any synaptic and cellular node diminished learning about cues. Our work provides a new perspective on how MSN subtypes interact to facilitate learning about aversive cues associated with foot shock.

## Results

### Neurokinin 1 receptor signaling is required for cue-dependent aversive learning

Since substance P is released by strongly salient stimuli ^23,24^ and is involved in many of the same aversive outcomes as D2-MSNs activation, we first tested whether putative substance P release and binding to its primary receptor, the neurokinin 1 (NK1) receptor, was necessary for aversive learning. A Pavlovian fear conditioning model was used as a model of aversive learning, where freezing was the main measure of learning (**Figure 1A**). To determine if substance P was required for associative aversive learning, mice were injected (i.p.) with the brain penetrant neurokinin 1 receptor antagonist L-733,060 (10 mg/kg) prior to fear conditioning. NK1 receptor antagonism suppressed cue-dependent, but not context dependent freezing (**Figure 1B**), without affecting locomotion (**Figure S1**). Intraperitoneal injection of L-733,060 prior to cue recall had no effect on cue recall (**Figure 1C**). These results suggest substance P signaling during aversive cue conditioning is required for the full expression of fear learning.

**Figure 1.**
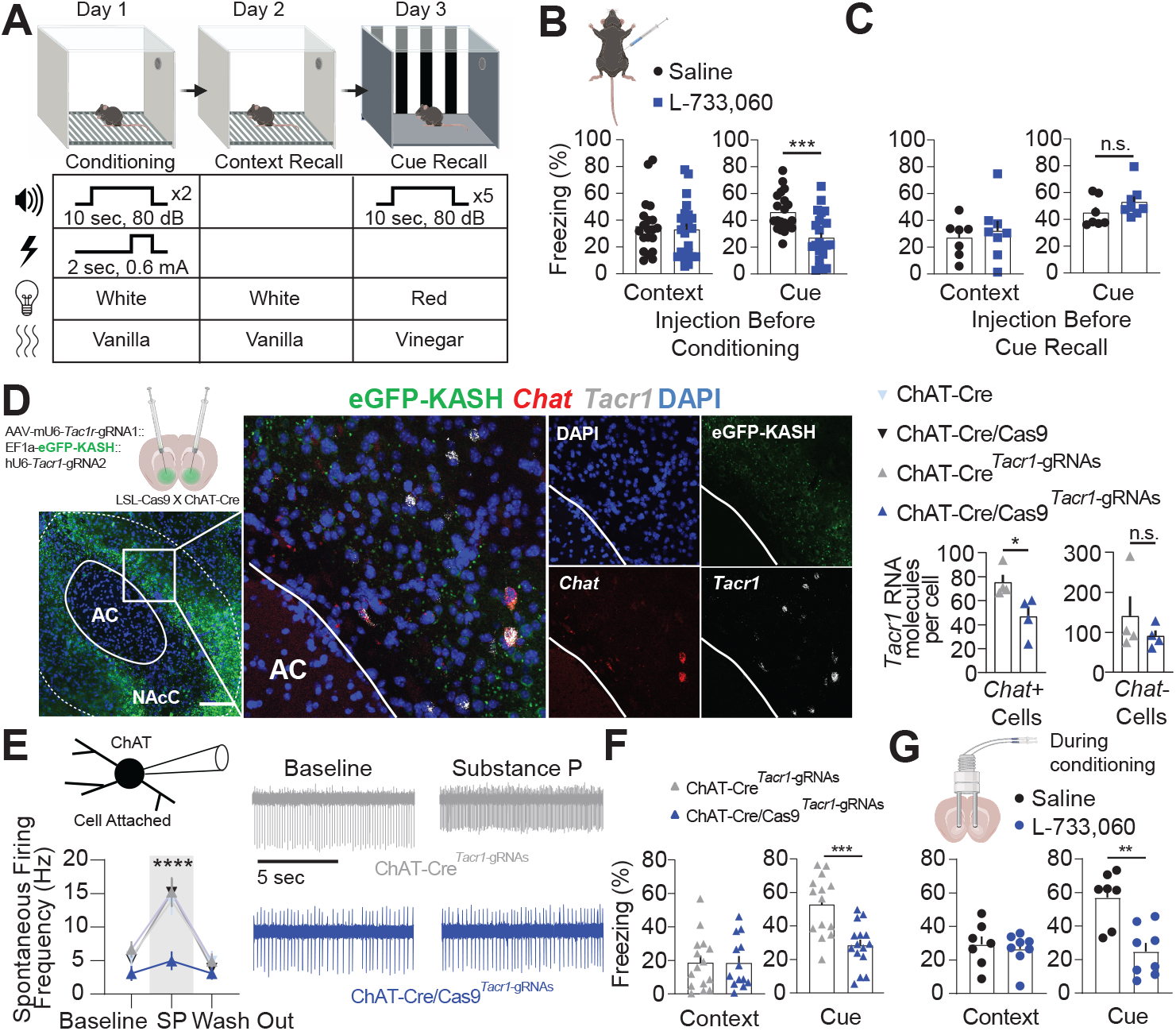
NAc substance P signaling is required for cue-dependent aversive learning. **(A)** Fear conditioning paradigm for aversive learning. **(B)** Injection of the NK1 receptor antagonist L-733,060 (10 mg/kg) or saline was given prior to conditioning. NK1 receptor antagonist suppressed freezing in response to the cue during cue recall (*p* < 0.001) and no difference in freezing was observed in the context (*p* > 0.05) (n = 20, 22 mice). **(C)** L-733,060 prior to cue recall does not change the percent of time freezing to conditioned cues (*p* > 0.05) (n = 7, 8). **(D)** Fluorescent image of EGFP-KASH expression in the NAc core. Scale bar 100 µm. Insert: Expression of EGFP-KASH (green), *Chat* RNA (red), *Tacr1* RNA (grey/white), and DAPI (blue). CRISPR-Cas9 mediated knockout of *Tacr1* decreased measured *Tacr1* RNA puncta in *Chat* RNA positive cells (*p* < 0.001) but not *Chat* RNA negative cells (*p* > 0.05) (n = 4 mice per group). **(E)** Graph and representative traces showing Cas9 and *Tacr1-*gRNA was required to suppress substance P(SP)-dependent activation of ChAT cells (Interaction *p*<0.0001; post-hoc Baseline vs. SP ChAT-Cre/Cas9*^Tacr1^ p* > 0.05, Mice/cells(N/n) = 4/10; Baseline vs. SP ChAT-Cre*^Tacr1^*, ChAT-Cre/Cas9, or ChAT-Cre *p* < 0.0001) (N/n = 4/9, 4/8, 3/7). **(F)** *Tacr1* knockout decreased freezing to the cue (*p* < 0.001), but not the conditioning context (*p* > 0.05) (n = 14, 15 mice). **(G)** Local NAc infusion of L-733,060 during conditioning suppressed freezing to conditioned cues (*p* < 0.01) but not the conditioned context (*p* > 0.05) (n = 7, 8). Data are represented as mean ± SEM. For detailed statistics see **Table S1**. See also **Figure S1**.

The NK1 receptor is found throughout the brain, including in regions known to be associated with fear conditioning (e.g., amygdala and periaqueductal grey). In the NAc, the NK1 receptor (encoded by the *Tacr1* gene) is found in all choline acetyltransferase (ChAT) expressing cholinergic cells. Therefore, we used a clustered regularly interspaces short palindromic repeats (CRISPR) Cas9 system, generated an AAV expressing guide RNAs for the *Tacr1* gene (*Tacr1*-gRNAs), and injected it in the NAc core of ChAT-Cre/Cas9 mice, which express Cas9 exclusively ChAT cells. RNA scope *in situ* hybridization was used to assess the efficacy of the knockout. Since we were unable to separate ChAT cells expressing or not expressing the *Tacr1* guide RNAs (see Methods), all *Chat* expressing cells in the NAc core where GFP expression was observed were quantified for *Tacr1* expression. A significant reduction in the *Tacr1* transcript in *Chat* positive cells was observed and no difference was observed in *Tacr1* expression in *Chat* negative cells (**Figure 1D**). Additionally, a functional and near complete inhibition of substance P-induced increase in ChAT firing was observed in mice expressing Cas9 and the *Tacr1*-gRNAs (**Figure 1E**). Next, we tested the functional knockout in fear conditioning behavior. NK1 receptor knockout suppressed cue-dependent, but not context-dependent freezing (**Figure 1F**), suggesting the cue-dependent reduction in freezing due to NK1 receptor antagonism was indeed an effect selective for the NAc core. In agreement with this finding, local NAc core infusion of L-733,060 during conditioning was capable of suppressing cue-dependent but not context-dependent recall compared to saline infused mice (**Figure 1G**). Overall, these experiments indicate substance P signaling is required for the full expression of cue-dependent recall of aversion-associated cues.

### D1-MSNs are required for cue-dependent fear conditioning

The primary source of substance P in the striatum is D1-MSNs which exclusively express and release the peptide and we have demonstrated activation of D1-MSNs releases substance P ^22^. Since substance P signaling is required for aversive learning and substance P is released in the striatum in response to salient aversive stimuli ^23^, we predicted D1-MSN activity would increase in response to salient, aversive stimuli. To selectively assay activity of D1-MSNs, Dyn-Cre mice were injected with AAV-DIO-GCaMP7f and implanted with fibers in the NAc core for assessment of *in vivo* calcium activity with fiber photometry as a means of assessing correlated neural activity (**Figure 2A**). We first acquired signals in response to random interval, unpredictable foot shocks. D1-MSN calcium activity scaled with the magnitude of foot shock (**Figure 2B-C**) but did not significantly vary across shock trials (**Figure 2D**). This scaling was not affected by injection of L-733,060 (**Figure S2A**). Next, we assayed activity during conditioning and cue recall. During conditioning, foot shock but not the novel cue increased D1-MSN calcium activity (**Figure 2E**). D1-MSN calcium activity was not increased by the cue during cue recall (**Figure 2F**), despite an increase in freezing (**Figure S2B**). Additionally, individual freeze events were not associated with changes in D1-MSN calcium activity (**Figure S2C**). We then scaled foot shock intensity to determine if the magnitude of the foot shock controlled both the D1-MSN calcium response and subsequent learning. As expected, freezing to the associated cue increased with foot shock intensity (**Figure 2G**). Interestingly, the area under the curve (AUC) of the shock-induced calcium signal during conditioning correlated significantly with the percent freezing (**Figure 2H**). Taken together, these results indicate the strength of D1-MSN activity during conditioning may be necessary for scaling cue-dependent aversive learning and recall.

**Figure 2.**
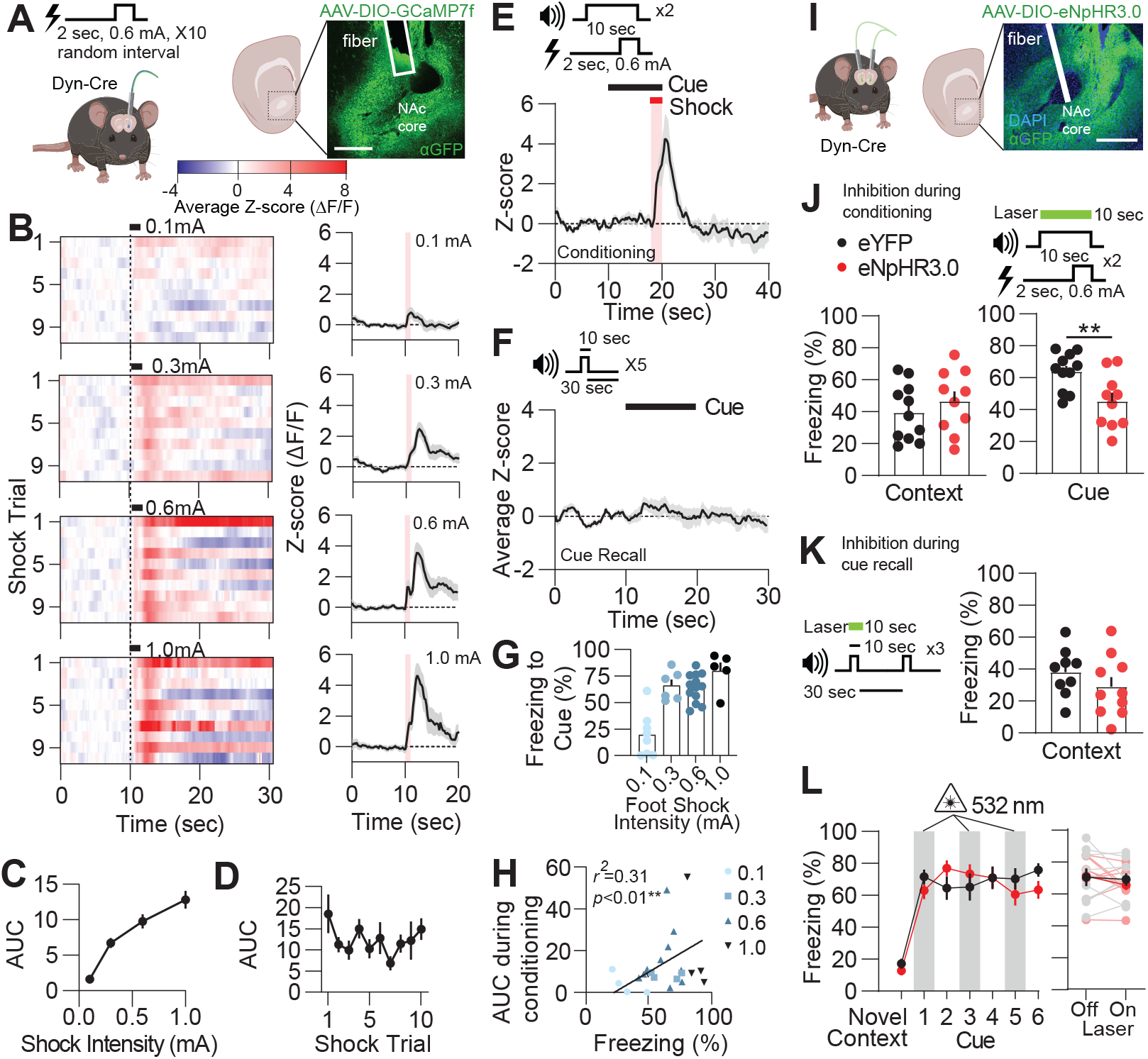
D1-MSN activity is required for cue-dependent conditioning. **(A)** Representation of a Dyn-Cre mouse injected with the calcium indicator GCaMP7f and a representative image of its expression in the NAc core with a fiber track. Scale bar 200 µm. **(B)** Heat maps and time plots representing average z-scores of D1-MSN calcium-driven fluorescence over increasing foot shock intensity (0.1-1.0 mA). **(C)** AUC from GCaMP7f fluorescent signals increased with shock intensity (Interaction *p* < 0.0001, n = 8-10 mice per shock intensity). **(D)** No difference was observed across individual shock trials at 0.6 mA (*p* > 0.05, n = 10 mice). **(E)** Averaged z-score D1-MSN calcium activity during fear conditioning. **(F)** Average z-score D1-MSN calcium activity in response to the condition cue during cue recall. **(G)** Conditioning foot shock intensity increasingly enhances freezing to the cue during recall (*p* < 0.0001). **(H)** AUC of the D1-MSN GCaMP7f signal significantly correlates with freezing to the cue during recall (*p* < 0.01) (n = 5-13 mice per group) **(I)** Expression of eNpHR3.0-eYFP, fiber track for optogenetic inhibition of the NAc core, and cartoon displaying bilateral implant and expression of the optogenetic construct in mice. **(J)** Inhibition of D1-MSNs during the cue in conditioning suppressed later cue recall (*p* < 0.01) but did not affect context recall (*p* > 0.05) (n = 11, 10 mice). **(K)** Halorhodopsin inhibition of D1-MSNs during cue recall had no effect on subsequent context recall. **(L)** Inhibition of D1-MSNs during cue recall had no effect on freezing (Laser off vs. on, *p* > 0.05) (n = 9, 10 mice). Data are represented as mean ± SEM. For detailed statistics see **Table S1**. See also **Figure S2**.

To test the necessity of D1-MSN activity during cue-dependent aversive learning, we used a temporally selective optogenetic strategy to inhibit D1-MSNs during cue-dependent behavior. Dyn-Cre mice were injected with AAV-DIO-eNpHR3.0 and implanted with optogenetic fibers in the NAc core (**Figure 2I**). Optogenetic inhibition of D1-MSNs did not affect locomotion (**Figure S2D-E**). D1-MSNs were inhibited selectively for 10 sec during the cue and shock during conditioning. D1-MSN inhibition during the cue and shock in conditioning diminished cue recall but not context recall (**Figure 2J**). In another group of mice, we selectively inhibited D1-MSNs during every other 10 sec cue to determine if D1-MSN inhibition reversibly inhibited cue-dependent freezing. No change in cue recall was observed from D1-MSN inhibition (**Figure 2K-L**). Altogether, these results indicate D1-MSN activity contributes to cue-dependent learning, but not recall.

### Substance P signaling is required for shock-driven acetylcholine release

Substance P is released from D1-MSNs in response to salient stimuli ^23,32^ and has a role in aversion ^33,34^. Additionally, substance P release promotes increased activity of ChAT neurons via the NK1 receptor ^22,35^. In addition, previous studies have indicated ChAT neuron exhibit phasic activity patterns in response to aversive stimuli *in vivo* ^26^. Therefore, it was predicted substance P would cause ChAT activity and acetylcholine release *in vivo*. An adeno-associated virus (AAV) containing the GPCR-activation-based sensor for acetylcholine (GRAB-ACh3.0) ^36^ was injected into the NAc core of mice and expressed in neurons (**Figure 3A**). *Ex vivo* slice studies demonstrated bath application of low (1 µM) and high (100 µM) concentrations of acetylcholine significantly increase bulk fluorescence intensity in a concentration dependent manner (**Figure S3A**). To evaluate acetylcholine binding *in vivo*, the virus was injected, a fiber was implanted in the NAc core, and fluorescent transients were observed with fiber photometry. In an open field chamber, mice were injected with the broad muscarinic antagonist scopolamine, which has been observed to block acetylcholine binding to the GRAB-ACh3.0 sensor ^36^. Scopolamine significantly diminished the average magnitude of the GRAB-ACh signal and suppressed the amplitude of all transients (**Figure S3B**). These results verify that the GRAB-ACh3.0 sensor causes fluorescence through acetylcholine release and binding to the GRAB-Ach3.0 receptor sensor.

**Figure 3.**
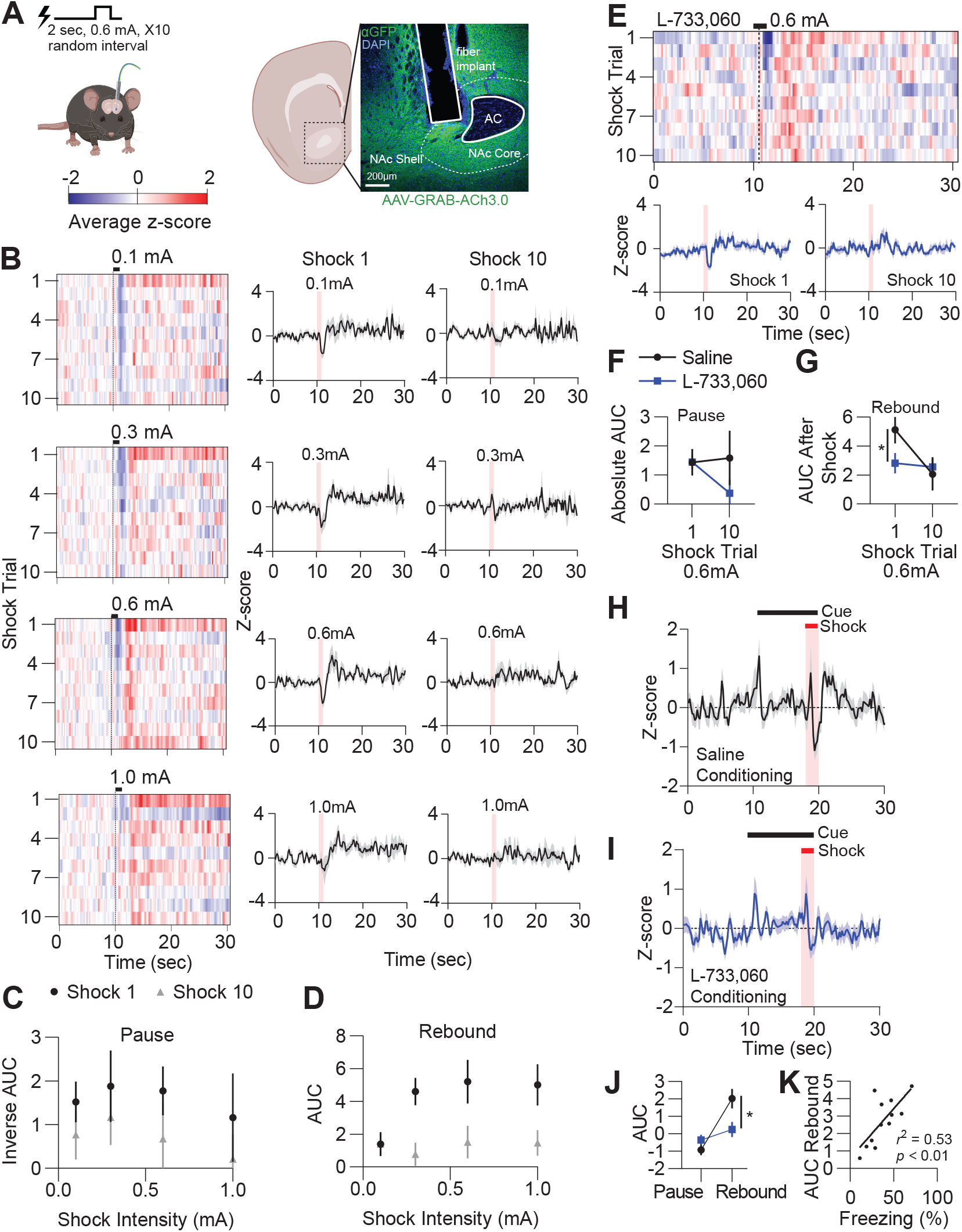
Acetylcholine release during foot shock is substance P-dependent. **(A)** Representative image of GRAB-ACh3.0 expression in the NAc core and cartoon of fiber placement. **(B)** Heatmaps and time courses of fluorescent GRAB-ACh3.0 signals from 10 random foot shocks at intensities of 0.1-1.0 mA amplitudes. Time courses are taken from average z-scores from shock 1 and shock 10. **(C and D)** AUC measurement of acetylcholine release at varying shock intensities for shock 1 and shock 10. **(C)** No differences were observed during the AUC (inverse to make positive) during pauses in acetylcholine-mediated GRAB-ACh3.0 fluorescence across shock intensities or repeated shocks (*p* > 0.05). **(D)**. Acetylcholine release varied across both AUC and shock intensity during the post shock rebound (Cue 1 vs. Cue 10, *p* < 0.05) (n = 9 mice). **(E)** Heat map and time course of GRAB-ACh3.0 fluorescent response after L-733,060 administration. **(F)** L-733,060 had no effect on the size of the pause across shocks (*p* > 0.05). **(G)** NK1 receptor antagonism suppressed AUC of the acetylcholine signal varied across both drug treatment (AUC Saline vs. L-733,060 during Cue 1, *p* < 0.05) and shock presentation (Cue 1 vs. Cue 10, *p* < 0.05) (n = 5, 6 mice). **(H-I)** Phasic changes in GRAB-ACh3.0 fluorescence during aversive cue conditioning in saline and L-733,060 treated mice. **(J)** L-733,060 significantly decreases the rebound AUC GRAB-ACh3.0 fluorescence during cue conditioning (p < 0.05, n = 8, 12). **(K)** The AUC during the rebound in conditioning significantly correlated with percent freezing to the cue during recall (*p* < 0.01, n = 12 mice). Data are represented as mean ± SEM. For detailed statistics see **Table S1**. See also **Figure S3**.

Next, to determine if acetylcholine is released by aversive stimuli in a substance P-dependent manner, foot shocks (10 total) were given at random, unpredictable intervals in behavior naïve mice while recording GRAB-ACh3.0 fluorescent activity in the NAc core. Shock alone produced a decrease in fluorescence (“pause”) followed by an increase (“rebound”) in acetylcholine-mediated fluorescence (**Figure 3B**), suggesting acetylcholine is released in a multi-phasic pattern. This phasic pause and rebound has been observed in rewarding stimuli in other studies ^46^. Similar to D1-MSN calcium activity, increasing the intensity of the shock increased the amount of rebound acetylcholine fluorescence following the shock without significantly changing the AUC of the pause, despite habituation to repeated shock (**Figure 3C-D**). Unlike D1-MSNs, this acetylcholine release habituated to the shock over repeated presentations (**Figure 3C-D**). Next to determine if the NK1 receptor was responsible for the rebound increase in acetylcholine fluorescence, L-733,060 was injected (*i.p.*) and mice were exposed to random foot shocks at 0.6 mA. Antagonism of the NK1 receptor attenuated the increase in acetylcholine-mediated fluorescence AUC during the rebound only (**Figure 3E-G**) and did not diminish habituation to the shock in the rebound condition (**Figure 3G**). These results suggest aversive stimuli increase acetylcholine release via substance P after foot shock and acetylcholine release scales with the intensity of an aversive stimulus.

To determine how aversive conditioned stimuli affect this signal, mice injected with GRAB-ACh3.0 in the NAc core were run through fear conditioning. Three prominent phasic events were observed in acetylcholine-mediated fluorescence: a response to the novel cue, a pause, and a rebound (**Figure 3H**). Like in the condition of the shock alone, the rebound response was attenuating (**Figure S3C**). The rebound fluorescence, but not the other features, was significantly diminished in mice treated with L-733,060 compared with saline injected mice (**Figure 3I-J**). During cue recall, acetylcholine fluorescence was only increased at the termination of the cue in saline treated mice but was increased at the start of the cue and the termination of the cue in L-733,060 treated mice (**Figure S3C-D**). This increase in acetylcholine fluorescence observed in L-733,060 treated mice is attenuated after the first presentation of the cue during cue recall (**Figure S3D**). This may suggest that the cue itself still appears to be novel to the mice with NK1 receptor antagonism. Interestingly, the rebound AUC significantly correlated with the time spent freezing to the cue during cue recall (**Figure 3K**). Overall, these results indicate that the increase in fluorescence following the shock is important for learning about cues predicting aversive foot shock.

### Shock-induced acetylcholine is important for learning about a conditioned aversive cue

To determine the necessity of NAc core ChAT cells are influencing learning about cues predicting foot shocks, we injected ChAT-Cre mice with an AAV-DIO-eNpHR3.0 or AAV-DIO-eYFP construct and inhibited ChAT neurons at two different time points during fear conditioning to align inhibition to time windows of GRAB-ACh fluorescence: during the conditioning cue or immediately after the foot shock (**Figure 4A**). Inhibition of ChAT cells had no observed effects on locomotion (**Figure S4**). Mice that received inhibition during the cue showed no change in context or cue recall compared to eYFP controls (**Figure 4B**). However, 5 sec of inhibition of ChAT cells following the shock, during the observed rebound, significantly diminished freezing to cues predicting shocks (**Figure 4C**), suggesting acetylcholine from ChAT neurons during the rebound condition is necessary for the full expression of aversive cue recall.

**Figure 4.**
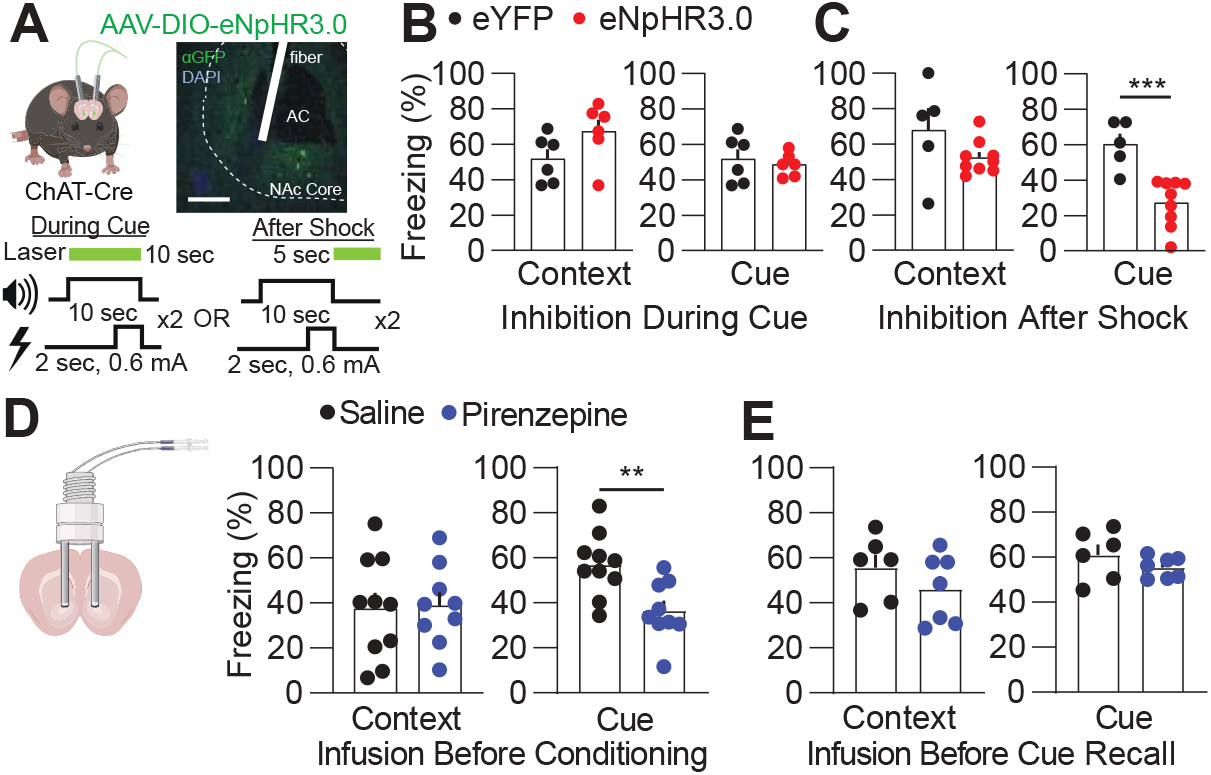
ChAT-mediated cholinergic signaling is required for cue-dependent aversive learning. **(A)** Expression of halorhodopsin in ChAT cells in the NAc core. Schematic showing inhibition paradigms: during the cue in conditioning or after the shock and cue during conditioning. **(B)** ChAT inhibition during the cue had no effect on context or cue recall (*p* > 0.05, n = 6 mice per group). **(C)** Inhibition of ChAT cells following the cue and shock during conditioning significantly decreased freezing to the cue during recall (*p* < 0.001), but did not affect context recall (*p* > 0.05) (n = 5, 9). **(D)** Selective infusion of the muscarinic 1 receptor antagonist pirenzepine (bilateral) prior to conditioning suppressed cue-dependent (*p* < 0.01) but not conditioning context dependent freezing (*p* > 0.05) (n = 10, 9). **(E)** Infusion of pirenzepine prior to cue recall had no effect on freezing (*p* > 0.05) in mice showing similar context recall (n = 6, 7). Data are represented as mean ± SEM. For detailed statistics see **Table S1**. See also Figure S4.

Substance P causes long-term potentiation (LTP) of excitatory transmission on D2-MSNs through enhanced acetylcholine release and activation of the muscarinic 1 receptor signaling pathway ^22^. This mechanism would provide a cell-type specific mechanism for enhancing activity of D2-MSNs to either promote aversive outcomes or to facilitate learning about aversive stimuli. To first determine if aversive conditioning and subsequent acetylcholine release acts through muscarinic 1 receptors on D2-MSNs, mice were implanted with cannulas in the NAc core and infused with the muscarinic 1 receptor-like antagonist pirenzepine (4 ng/µL, bilateral) prior to conditioning or prior to cue retrieval. Compared to saline infused controls, cue recall was suppressed when pirenzepine was infused prior to conditioning (**Figure 4D**), but not prior to cue recall (**Figure 4E**).

### Substance P promotes plasticity on D2-MSNs during cue-dependent aversive learning

D2-MSNs are likely to receive this excitatory muscarinic 1 receptor signaling more strongly as D1-MSNs also express the inhibitory muscarinic 4 receptor which could suppress this signaling. To determine if muscarinic 1 receptor plasticity occurred on D2-MSNs in a substance P-dependent manner, we conditioned mice to a cue after injection with saline or L-733,060 and conducted whole-cell electrophysiology recordings 1 day after cue recall. Substance P-dependent LTP on D2-MSNs is primarily induced through insertion of GluR2-lacking AMPA receptors ^22^. These GluR2-lacking AMPA receptors are sensitive to intracellular peptides at positive membrane potentials and excitatory synaptic currents at positive potentials show inward rectification when these receptors have been inserted in the synaptic membrane. Therefore, we assessed inward rectification of electrically evoked excitatory post-synaptic currents (EPSCs), which would be suggestive of increased membrane insertion of GluR2-lacking AMPA receptors. Mice that received conditioning (Cue + Shock) displayed significant rectification at +40 mW as compared to non-conditioned mice (Cue only) (**Figure 5A**). Additionally, mice injected with L-733,060 displayed no difference in rectification compared to non-conditioned mice, suggesting that suppressing substance P signaling was responsible for LTP observed on D2-MSNs (**Figure 5A**). As a secondary measure, we compared sensitivity of electrically evoked currents on D2-MSNs from conditioned and non-conditioned mice to the selective GluR2-lacking AMPA receptor antagonist 1-naphthyl acetyl spermine trihydrochloride (NASPM) (100 µM). NASPM significantly diminished currents in conditioned mice relative to non-conditioned mice, verifying the presence of more GluR2-lacking AMPA receptors (**Figure 5B**). Furthermore, amplitude, but not frequency of spontaneous excitatory post-synaptic currents (sEPSCs) was increased on D2-MSNs (**Figure 5C**). Shock only mice or context only mice displayed no changes in inward rectification on D2-MSNs (**Figure S4**). D1-MSNs did not display inward rectification from any groups assayed (**Figure 5D and S4**). Results indicate signatures of substance P-dependent LTP are expressed selectively in D2-MSNs following cued aversive learning.

**Figure 5.**
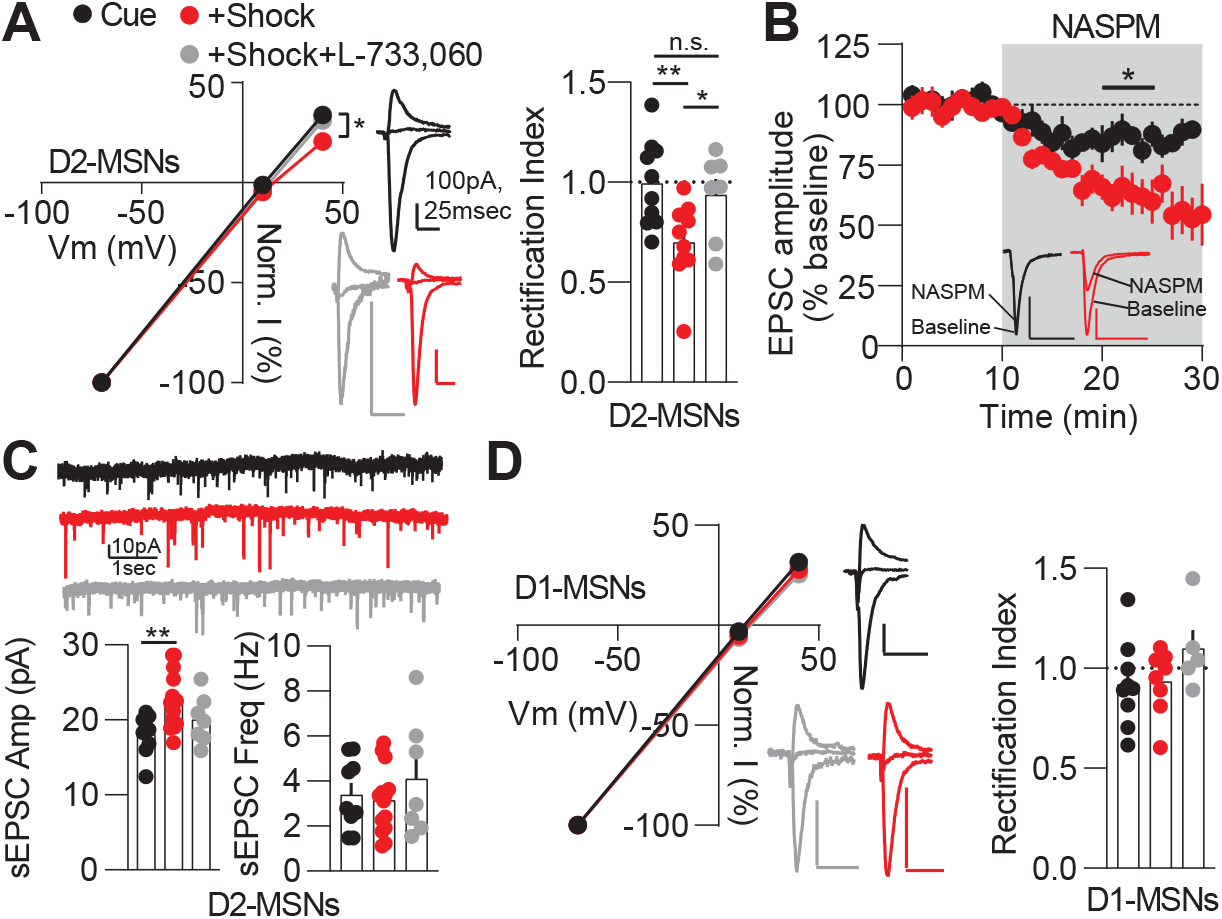
Excitatory plasticity on D2-MSNs is caused by cue-dependent aversive learning. **(A)** Cue/shock conditioning enhanced inward rectification of the I-V curve generated from electrically evoke excitatory currents on D2-MSNs (Interaction *p* < 0.001) from the cue/shock group compared to cue only (post-hoc *p* < 0.05) and cue/shock with L-733,060 (post-hoc *p* < 0.05). Rectification index indicated more rectification for D2-MSNs from cue/shock mice (Interaction, *p* < 0.01) compared to cue only (post-hoc, *p* < 0.01) and cue/shock with L-733,060 (post-hoc, *p* < 0.05) (N/n (Mice/cells) = 5/10, 5/10, 4/7). **(B)** NASPM suppressed currents more from D2-MSNs from cue/shock mice compared to cue only mice (25-30 min average, *p* < 0.05) (N/n = 4/6, 4/5). **(C)** Spontaneous excitatory post-synaptic current (sEPSC) amplitude (*p* < 0.05), but not frequency (*p* > 0.05) is increased in D2-MSNs following fear conditioning (N/n = 6/14, 5/9, 4/7). **(D)** D1-MSNs do not show inward rectification across conditions (*p* > 0.05) (N/n = 6/8, 4/9, 4/5). Data are represented as mean ± SEM. For detailed statistics see **Table S1**. See also **Figure S5**.

### D2-MSNs activity is necessary for recall of aversive-conditioned cues

If LTP is selectively expressed on D2-MSNs and is dependent on aversive cue associations, then it is likely D2-MSNs are active in response to aversively conditioned cues. To verify this activity in real time and *in vivo*, A2a-cre mice were injected with AAV-DIO-GCaMP7f in the NAc core and implanted with fibers for fiber photometry, A2a-Cre mice were injected with AAV-DIO-GCaMP7f and implanted with fibers for photometry (**Figure 6A**) and subjected to 10 unpredictable foot shocks at various foot shock intensities. D2-MSNs responded similarly to all foot shock intensities (**Figure 6B**) and L-733,060 did not alter D2-MSN calcium response to foot shock (**Figure S6A**). In contrast to D1-MSNs, D2-MSN calcium activity did not increase with increasing foot shock intensity (**Figure 6C**) and responses were variable across shock events (**Figure 6D**). Like D1-MSNs, D2-MSNs responded to foot shock during conditioning, but not the cue (**Figure 6E**). Unlike D1-MSNs, D2-MSNs were activated during cue recall (**Figure 6F**) in response to the associated aversive cue and this signal did not habituate over repeated plays of the cue (**Figure S6B**). Furthermore, D2-MSN calcium activity did not increase in response to individual freezing events, suggesting freezing alone did not explain D2-MSN calcium signals (**Figure S6C**). To determine if NK1 receptors were required during conditioning to cause this increase to the cue, L-733,060 was injected in mice prior to conditioning. NK1 receptor antagonism did not affect D2-MSN calcium response to the shock during conditioning (**Figure 6G**). Next, mice conditioned in the presence of L-733,060 were exposed to conditioned cues. Photometry mice injected with L-733,060 showed significantly reduced freezing to the conditioned cue compared to saline injected mice (**Figure S6D**). D2-MSN calcium activity did not increase in response to conditioned cues in L-733,060 treated mice (**Figure 6H-I**). In addition, AUC during the cue correlated with freezing (**Figure 6J**). Taken together, this suggests NK1 receptors are required for D2-MSN activity in response to conditioned aversive cues.

**Figure 6.**
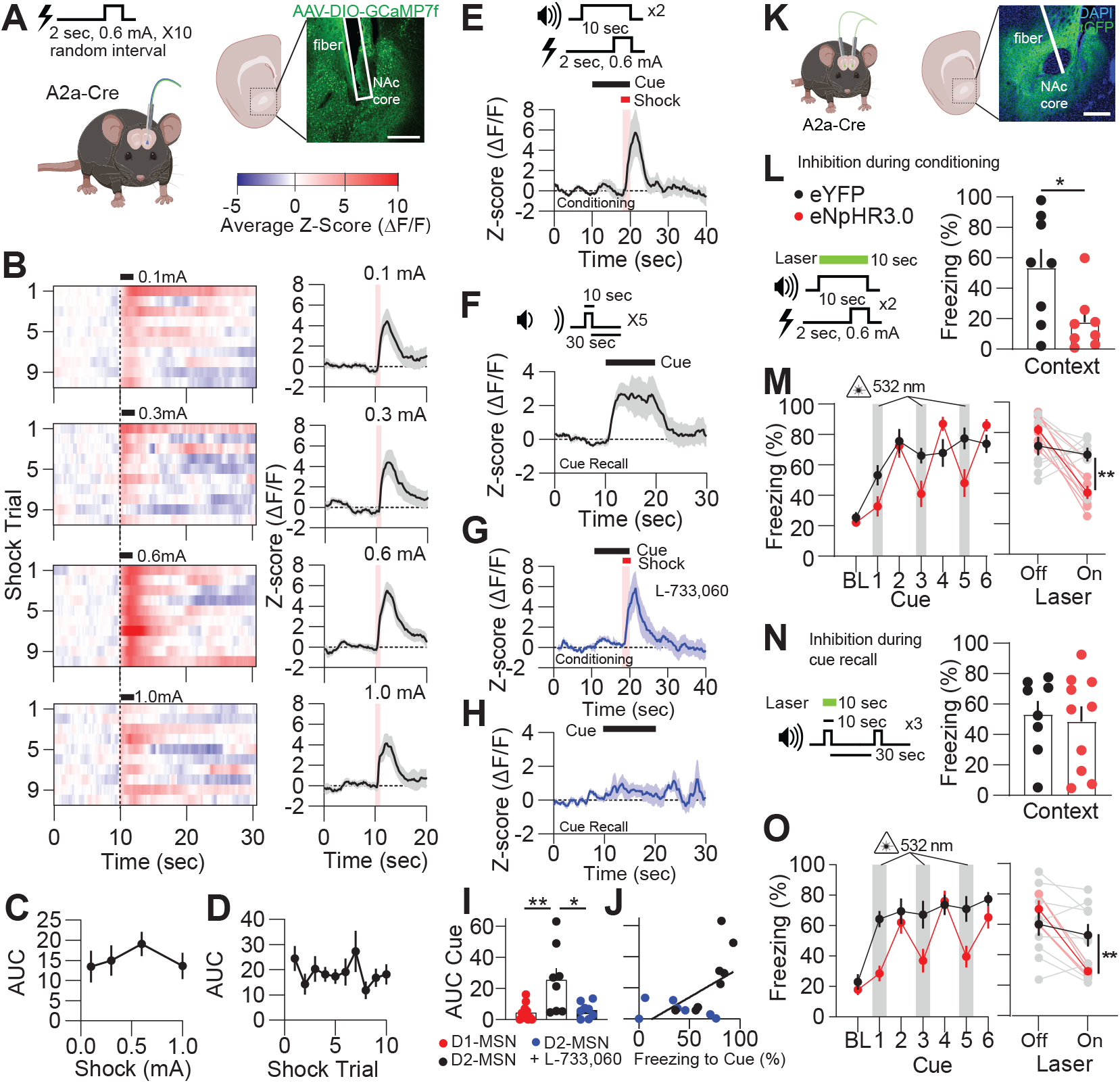
D2-MSNs are required for cue-dependent aversive learning. **(A)** A cartoon of an A2a-Cre mouse injected with the calcium indicator GCaMP7f and a representative image of its expression in the NAc core with a fiber track. Scale bar 200 µm. **(B)** Heat maps and time plots representing average z-scores of D2-MSN calcium-driven fluorescence over increasing foot shock intensity (0.1-1.0 mA). **(C)** AUC of an average of all 10 foot shocks indicated was no change in response across foot shock intensity (*p* > 0.05). **(D)** AUC fluorescence from D2-MSN GCaMP7f does not change across all foot shock trials at 0.6 mA intensity (*p* > 0.05,) (n = 10 mice). **(E)** D2-MSNs responded to the first conditioning shock, but not the cue. **(F)** Following conditioning, calcium signals from D2-MSNs were responsive to the conditioned cue. **(G)** D2-MSN calcium signals responded to the foot shock during conditioning after being injected with L-733,060. **(H)** Cue-driven calcium transients from D2-MSNs were not observed in mice injected with L-733,060. **(I)** AUC fluorescent transients during cue recall from mice expressing GCaMP7f in D2-MSNs were larger than in mice expressing GCaMP7f in D1-MSNs (*p* < 0.01) and in mice injected with L-733,060 expressing GCaMP7f in D2-MSNs (*p* < 0.05) (n = 11, 8, 8).**(J)** AUC during the cue significantly correlated with freezing to the cue (*p* < 0.05, n = 8, 8). **(K)** eNpHR3.0-eYFP expression in D2-MSNs within the NAc core and cartoon depicting bilateral inhibition of the NAc core. **(L)** Inhibition of D2-MSNs during the cue in conditioning decreased freezing during context recall (*p* < 0.05). **(M)** Inhibition of D2-MSNs reversibly suppressed freezing to recalled cues (Over cues *p* < 0.01; Laser off vs on *p* < 0.01; post-hoc laser on eYFP vs. eNpHR3.0 *p* < 0.01) (n = 8, 8). **(N-O)** Inhibition of D2-MSNs reversibly suppressed freezing to recalled cues (Over cues *p* < 0.01; Laser off vs on *p* < 0.0001; post-hoc laser on eYFP vs. eNpHR3.0 *p* < 0.05) in mice that showed no difference in context recall (*p* < 0.05) (n = 9, 6 mice). Data are represented as mean ± SEM. For detailed statistics see **Table S1**. See also **Figure S6**.

Next, to determine if D2-MSN activity was required for recall of associated cues, AAV-DIO-eNpHR3.0 was injected, and fibers were implanted into the NAc of A2a-cre mice for selective optogenetic inhibition of D2-MSNs (**Figure 6K**). To compare the necessity of D2-MSNs to D1-MSNs in conditioning, mice received optogenetic inhibition of D2-MSNs during both cues during conditioning (**Figure 6K**). Surprisingly, inhibition of D2-MSNs during the cue in conditioning significantly diminished freezing in the context (**Figure 6L**). As expected, D2-MSN inhibition during conditioning had no effect on freezing to cues during cue recall when D2-MSNs were not inhibited (**Figure 6M**). Despite this, inhibition of D2-MSNs during cue recall diminished freezing (**Figure 6M**), suggesting D2-MSNs are required for the full expression of freezing to aversively conditioned cues. To test this outcome without the potential confounds of inhibition during cue conditioning, D2-MSNs were inhibited during the 10 second cue on every other cue (**Figure 6N**). D2-MSN inhibition reversibly inhibited freezing to the cue (**Figure 6O**) without affecting freezing to the context (**Figure 6N**). This suppression in freezing was not dependent on locomotor effects (**Figure S6E-F**). Taken together, associative aversive stimuli promote activity of D2-MSNs in response to associative cues and this response is required expression of cue-dependent aversive learning.

## Discussion

Our work has demonstrated a di-synaptic, local circuit within the NAc core that is critical for processing aversive cues, with necessary roles for NAc core D1-MSN and D2-MSN activity at different stages of learning and recall (**Figure 7**). While the activity does not appear to be required for the full expression of the fear behavior, activity of these subtypes significantly modulates the overall expression of the cue-dependent behavior, suggesting it does indeed play a critical role in learning. We have shown that D1-MSN activity and substance P is required for learning during cue-dependent aversive conditioning and D2-MSNs are required for recall of conditioned cues. This D2-MSN activity change is likely due to excitatory LTP ^22^ produced through muscarinic receptor activation, which would enhance the likelihood of D2-MSN activation during excitatory input that encodes salient stimuli. Overall, our results provide evidence for the involvement of the Nucleus Accumbens core in Pavlovian learning that might support later motivated output.

**Figure 7.**
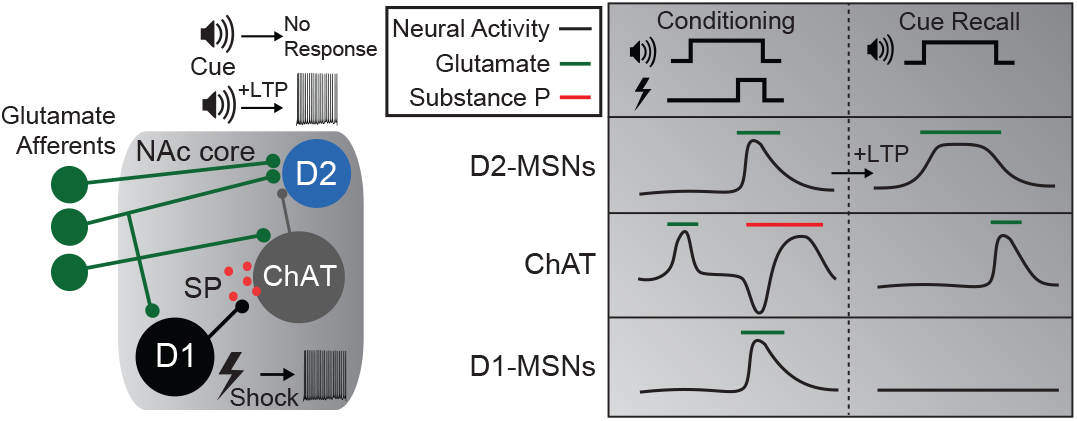
Model for the role of the NAc core in cue-dependent fear learning. NAc circuitry involved in aversive cue learning (left). Shock activates D1-MSNs releasing substance P and activating ChAT neurons. ChAT neurons release acetylcholine selectively during cue/shock pairings and promotes glutamatergic long-term potentiation (LTP) on D2-MSNs rendering them more responsive to the cue. Timing of putative activity and release of neurotransmitters from NAc neuronal subtypes during cue conditioning and recall (right).

This work is consistent with previous studies showing MSNs respond to different aspects of associative reward conditions, where D1-MSNs respond to outcomes and D2-MSNs respond to associative cues ^21^. We found similar outcomes in response to an aversive stimulus, where D2-MSNs were active in response to a learned cue but not to novel cues and D1-MSNs were active in response to the shock alone. Together these findings indicate MSN subtypes are responding to salient events, including rewarding events, further supporting the hypothesis that the NAc core supports learning about salient cues ^1^. Furthermore, the duration of MSN subtype responses is temporally different in both rewarding ^21^ and aversive-associated stimuli. D1-MSNs rapidly respond to the non-associative, salient stimuli and D2-MSNs tend to display extended activity following associative cue presentation. We speculate D2-MSNs may be actively engaged in either retaining expression of the learned association or suppressing additional associations. More work will be needed to test the temporal nature of D1-MSN and D2-MSN activity in response to associative conditioning and learning.

We provide a role for D1-MSNs and the neuropeptide substance P in mediating aversive learning. D1-MSNs are thought to facilitate reward; however, the data presented here and data from others assessing activity in the NAc core ^38,39^ and NAc shell ^40,41^ suggest aversive stimuli activate D1-MSNs. Our data indicates this activity participates in associative learning through the release of peptides, activation of cholinergic neurons, and release of acetylcholine. Interestingly, scaling up D1-MSN activity appears to scale acetylcholine release, which is consistent with increasing substance P levels ^22^. While acetylcholine may act as an attentional signal facilitating the salience of associative stimuli ^42,43^, we have discovered substance P works upstream to promote ChAT activation, acetylcholine release, and plasticity on D2-MSNs, which may be dependent on the concentration of neuromodulator release in response to scaled stress. This plasticity facilitates information transfer from excitatory brain regions to the NAc core. As the effect of substance P and acetylcholine plasticity is cue-dependent but not context-dependent, it is likely that extensive excitatory projections arising from limbic regions that are involved in Pavlovian conditioning including the basolateral amygdala ^44,45^ or other regions involved in aversion such as the paraventricular thalamus ^47^ may facilitate this D2-MSN activity during cue recall. In contrast, strong projections to D1-MSNs from the ventral hippocampus ^40,49^ may facilitate D1-MSN activity for substance P release. Future experiments will need to assess what inputs undergo substance P, acetylcholine-dependent potentiation.

Our work shows acetylcholine release plays a significant role in aversive learning and its release is likely driven by substance P/NK1 receptor signaling. This could result from varying the upstream activity of D1-MSNs, subsequent substance P release, and the activation of ChAT neurons. Our previous work has shown that the activity of ChAT cells can be graded by varying concentration of substance P ^22^. We demonstrated D1-MSN activity increases with foot shock intensity, likely leading to increasing putative substance P release, and this might be a method of grading acetylcholine release and learning or response to an aversive stimulus. Other factors are likely contributing to this effect as well. We cannot rule out the role of dopamine input via D2 receptors ^46^, midbrain GABA input ^48^, thalamic input ^50^, intrinsic inhibition ^51^ or other mechanisms in generation of these cue-dependent outcomes. In fact, substance P and ChAT acetylcholine is known to promote dopamine release ^52–55^. Dopamine release could accompany the increase in acetylcholine observed in response to novelty ^56^ or initial phasic transients of foot shock. Therefore, selective plasticity could occur due to coordinated acetylcholine and dopamine release during learning.

Despite a non-habituating D1-MSN calcium signal following repeated foot shocks, the observed acetylcholine signal habituated to repeated foot shocks. After binding of substance P to the NK1 receptor and activation, rapid beta-arrestin-mediated endocytosis occurs ^57,58^ and attenuates additional responses to substance P for 5-15 min ^22,35,57^. At this point, NK1 receptor activation of ChAT cells would be less sensitive to substance P, thereby decreasing the size of the signal. Therefore, since we observed the acetylcholine release to be NK1 receptor dependent, we predict the habituation of the signal from shock 1 to shock 10 is due to NK1 receptor internalization. This, along with the NK1 receptor antagonist attenuation of the acetylcholine signal, indicates the increase in acetylcholine release is most likely due to release of substance P and activation of the NK1 receptor on ChAT neurons. Such a mechanism may suppress the likelihood of additional stimuli promoting plasticity that underlies associative learning. In addition, the D2-MSN foot shock signal is also rapidly attenuated and much smaller after repeated shocks. Mice only required two cue shock pairings for associating, perhaps indicating the plasticity on D2-MSNs occurs quickly, within the first few associations. This may further suggest that particularly salient stimuli can produce this plasticity rapidly.

One curious aspect of the findings is the activity observed in D1-MSNs, D2-MSNs, and acetylcholine is induced by foot shock alone. It is likely the release of substance P and subsequently acetylcholine interacts with other input on D2-MSNs to promote plasticity. This may include activity from glutamatergic inputs and ventral tegmental area dopamine. In fact, some evidence suggests the pause in cholinergic neurons is partially mediated by dopamine input to the striatum ^46^ and NAc D2-MSN spine dynamics in response to learned cues is dependent on dopamine ^16,51^. Furthermore, coordinated pre-synaptic and post-synaptic activity via glutamatergic input and muscarinic 1 receptor signaling on D2-MSNs may provide a window for spike timing dependent plasticity. These secondary sources may also explain the difference in the role of D2-MSNs in contextual and cue recall, as inhibition of D2-MSNs during conditioning suppressed context recall as well. These other sources will need to be investigated in future studies to fully understand the mechanisms that drive these outcomes.

Overall, we demonstrate substance P signaling and plasticity in the NAc core is critical for learning about salient, aversive cues. We uncovered a new role of D1-MSN local communication to facilitate acetylcholine plasticity on D2-MSNs that is relevant to cue-dependent aversive learning. Enhanced processing of salient stimuli, through this substance P, D2-MSN mechanism could lead to exacerbated response to aversive stimuli and could enhance susceptibility to aversive stimuli that cause stress or anxiety disorders. Therefore, targeting these neurobiological mechanisms may be useful in therapies aiming to treat anxiety and mood disorders.

## Acknowledgements

We would like to thank Dr. Geoffrey Schoenbaum for his assistance in conceptualization of the fear conditioning and latent inhibition experiments. This work was supported by that National Institutes of Mental Health R00MH123673, University of South Carolina College of Pharmacy, and the National Institute on Drug Abuse Intramural Research Program. Construction and packaging of the AAV vector expressing mTACR1 gRNAs was performed by the National Institute on Drug Abuse Genetic Engineering and Viral Vector Core, RRID:SCR_022969.

## Author contributions

Conceptualization, T.C.F.; Methodology and Investigation, A.B., C.G., C.S., C.R., D.B., E.M., A.S., H.H., and T.C.F.; Writing, T.C.F.; Review and Editing, A.B., C.G., A.S., C.R., and T.C.F.; Supervision, T.C.F.

## STAR Methods

### RESOURCE AVAILABILITY

#### Lead contact

Further information and requests or resources and reagents should be directed to and will be fulfilled by the lead contact, T. Chase Francis.

#### Materials availability

Plasmids generated in this study have been deposited to Addgene (pOTTC1553, Cat #: 131682; pOTTC1642, Cat #: 195018) and are publicly available as of the date of publication. All data reported in this paper will be shared by the lead contact upon request and can be found on the Open Science Framework platform (DOI 10.17605/OSF.IO/4FE3M).

#### Data and code availability

Python code generated to extract calcium and acetylcholine events is available through Github.com (https://github.com/FrancisLab-Repository/Photometry-Event-Extract.git). In some experiments, ezTrack was used to quantify freezing behavior or locomotion ^61^.

### EXPERIMENTAL MODEL AND SUBJECT DETAILS

#### Animals

Female and male mice 10-15 weeks old were used in all studies. For GRAB-ACh3.0 virus, cannula infusion experiments, and systemic drug injections without imaging or electrophysiology, C57Bl6/J mice (JAX: 000664) were used. For optogenetic inhibition experiments either B6;129S-Pdyntm1.1(cre)Mjkr/LowlJ (Dyn-Cre; JAX: 027958) or Tg(Adora2a-cre)KG139Gsat/Mmucd (A2a-Cre; MMRRC: 031168-UCD). For CRISPR-Cas9 experiments, B6;129-Gt(ROSA)26Sortm1(CAG-cas9*,-EGFP)Fezh/J (LSL-Cas9; JAX: 024857) mice were crossed with B6;129S6-Chattm2(cre)Lowl/J (ChAT-Cre) mice for conditional expression of Cas9 in ChAT-Cre cells. All transgenic animals were bred in house. Mice were fed *ad libitum* and were housed in a reverse light/dark (12/12 hr). Experimental procedures were approved and conducted in accordance with NIDA-IRP ACUC, University of South Carolina IACUC, and NIH guidelines.

### METHOD DETAILS

#### Surgical methods

Mice used in fluorescence imaging, optogenetic, or CRISPR knockdown experiments were anesthetized with ketamine/xylazine for intracranial AAV injection, hair was removed, scalps were cleaned with ethanol and betadine, and lidocaine was used prior to scalp incision. Body temperature was maintained with an electric heating pad. Mice received bilateral stereotactic injections (0.3-0.5 µL) of either AAV9-hsyn-GRAB-ACh3.0 (Addgene; plasmid# 121922; packaged by Vigene Biosciences)^36^, AAV9-syn-FLEX-jGCaMP7f (Addgene; 104492-AAV9), AAV5-EF1a-DIO-eNpHR3.0-eYFP (Addgene; 26966-AAV5), or AAV1-mU6-Tac1r-gRNA1::EF1a-NucEnv-EGFP::hU6-Tacr1-gRNA2 in the NAc core (created and packaged by NIDA GEVVC Core) (A/P: 1.6, Lat: 1.6, D/V: -4.4, 10° angle). For photometry or optogenetics, mice were then implanted with either a 400 µm photometry fiber (Doric) or 200 µm core optogenetic fiber fixed with head screws and dental cement. Following surgery, mice were injected with warm saline to replenish fluids and given carprofen (5 mg/kg) for pain relief during recovery.

#### *In vivo* fiber photometry, *ex vivo* imaging, and *in vivo* optogenetics

*In vivo* fiber photometry was used to assess cell-type specific calcium activity (GCaMP7f) or acetylcholine release (GRAB-ACh3.0) using a Tucker-Davis Technologies LUX RZ10 fiber photometry system. Signals were tested and mice were habituated for 3-5 days prior behavioral recordings. For all recordings, sensors were excited at 470 nm to acquire the fluorescent emission signal and the isosbestic point for GFP at 405 nm as a normalization control. Excitation signals were generated by two light-emitting diodes and emission signals were collimated through a Doric fluorescent mini cube before being processed and digitized using the Synapse software suite. Analysis was performed offline using a custom Python script which extracted fluorescent signals, smoothed the signal with a linear fit to the 405 nm signal and applying the regression to the 470 nm signal, calculated ΔF/F, aligned signals based on a timestamp, extracted fluorescent signals around events, and normalized signals (z-score) based on the 10 seconds prior to an event. Area Under the Curve (AUC) was assessed to find mean changes in calcium or acetylcholine activity. The following intervals were used for AUC calculation in the described experiment: for foot shock during fear conditioning or during shock alone, 0-5 sec from the initiation of the shock; for cue recall, 0-10 sec from the start of the cue; for shock-related decrease in ACh-mediated fluorescence, 0.5-2 sec after from the initiation of the shock; for shock-related rebound in ACh-mediated fluorescence, 2.5 to 7 sec from the initiation of the shock.

For *ex vivo* imaging of GRAB-ACh3.0, slices were prepared as discussed in the electrophysiology methods section. Acetylcholine (1 or 100 µM) was bath applied after a 2 min baseline. Videos were acquired with a 40x objective at a 10 Hz frame rate using QCapturePro7 and analyzed offline with ImageJ. A subregion 100×100 µm devoid of fiber tracts in the NAc core was chosen in each image and analyzed for fluorescence intensity over time. Fluorescence traces were flattened using a linear regression fit and normalized (z-score) to the 1 min prior to acetylcholine bath application.

For optogenetic inhibition experiments, mice were habituated to patch cables 5 days prior to behavior. Percent transmission was calculated during creation of optogenetic fibers and 594 nm light was delivered at 3-5 mW from a DPSS laser (OptoEngine LLC). Optogenetic inhibition was delivered during the onset of the cue and one second after the offset of the cue for all conditioning cues and on every other cue during cue recall.

#### Behavior

Pavlovian fear conditioning was conducted over a 3 day period (**Figure 1A**) in Med Associates Fear conditioning chambers contained in sound attenuating cabinets. Prior to each phase of fear conditioning, mice were habituated in a separate room for at least 1 hour in the lighting condition of the test. Conditioning was conducted in white light with vanilla scent (10% vanilla extract in ethanol) chamber with a grid floor. Following a 2 min baseline, 2 white-noise (80 dB) cues co-terminating in a 0.6 mA foot shock separated by a 30 sec interval. The next day mice were exposed to the conditioning context for 5 min. On the third day, mice were placed in a new context where floors were covered with white, opaque plexiglass, a curved white plexiglass insert, no light (red light in behavior room), and vinegar scent (1% acetic acid in water). After a 2 min baseline, 5-6 cues at 30 sec intervals were played back. In all tests, freezing was measured using VideoFreeze (Med Associates), which has been thoroughly validated in relation to hand scoring ^59^. Freezing thresholds were set at least 30, 0.033 sec frames of frozen motion and motion indices below 6 for mice without fiber or optogenetic tethers and below 500 for mice with optical tethers. For mice exposed to foot shock only, after a 2 min baseline each mouse was exposed to an unpredictable foot shock set to random intervals at least 30 sec between each shock. Mice received one intensity of foot shock per day, and each subsequent day were subjected to increasing foot shock intensities or decreasing foot shock intensities to control for previous foot shock intensities from interfering with the signal for any subsequent foot shock.

#### Electrophysiology

Whole-cell electrophysiological recordings were done one day following cue recall. Mice were anesthetized with Euthasol, brains were extracted, and cut (300 µM) in ice cold N-Methyl D-Glucamine (NMDG) cutting artificial cerebrospinal fluid (ACSF) solution (in mM): 92 NMDG, 20 HEPES, 25 glucose, 30 sodium bicarbonate, 1.2 sodium phosphate, 2.5 potassium chloride, 5 sodium ascorbate, 3 sodium pyruvate, 2 thiourea, 10 magnesium, 0.5 calcium chloride, osmolarity 305-310 mOsm, pH 7.34. Immediately after slicing, slices were placed in warm (32°C) NMDG solution for 2-5 min, then transferred to HEPES holding solution at room temperature which was the same as NMDG solution except NMDG was replaced with 92 mM sodium chloride and contained 1 mM magnesium chloride and 2 mM calcium chloride. Recording began 1 hr after brain slicing was complete and was performed at 33°C. Slices were perfused at a rate of 2 mL/min with ACSF (in mM): 125 sodium chloride, 26 sodium bicarbonate, and 11 glucose, 2.5 potassium chloride, 1.25 sodium phosphate, 2.4 calcium chloride, 1 magnesium chloride, osmolarity 305-310 mOsm, pH 7.34. To block inhibitory currents, synaptic recordings were done holding cells at -70 mV in the presence of picrotoxin (100 µM) and with a cesium methanesulfonate internal solution (in mM): 117 cesium methanesulfanate, 20 HEPES, 2.8 sodium chloride, 4 magnesium ATP, 0.3 sodium GTP, 0.4 EGTA, pH 7.30, osmolarity 280-287 mOsm. For IV curves, spermine (100 mM) was added to the pipette solution and currents were recorded in the presence of APV to isolate AMPA receptor currents. NASPM (100 µM) was used to isolate currents from inserted GluR2-lacking AMPA receptors. For cell-attached ChAT recordings, ACSF was used as the internal solution.

Whole-cell currents or cell attached currents were obtained with borosilicate glass pipettes (whole cell: 1.7-3.0 MΩ; cell attached: 3.0-4.0 MΩ) and signals were amplified and digitized using a Multiclamp 700B amplifier (4kHz low-pass filter for synaptic, 1kHz for cell-attached or spontaneous EPSCs) and Digidata 1322 digitizer (10 kHz), respectively (Molecular Devices). Synaptic currents were evoked with a concentric bipolar stimulating electrode (FHC) and evoked currents were stabilized for 5-10 min prior to data acquisition. Series resistance drifts of more than 20% led to exclusion of the cell from analysis. D1- and D2-MSNs were identified under the visual guidance of a BX61 Olympus microscope and Tdtomato fluorescence. Cholinergic neurons were identified by cell size (40-60 µm diameters) and high baseline firing rates.

#### CRISPR-Cas9 for *Tacr1* knockout

The SpCas9 guide RNA seed sequences within the first exon of mouse *TACR1* were identified and scored using the Guide Design Tool at CRISPR.mit.edu. Two gRNAs (gRNA 1 A55: CCGTATAGGCGGCTGCCCAA, and gRNA 2 B108: TTCCGTGGTGGGCAACGTAG) were chosen based on their likelihood for functioning as a “nickase-compatible” pair. These sequences were inserted into plasmids already containing RNA pol III promoters and SpCas9 gRNA scaffold sequences. (gifts from Charles Gersbach; Addgene plasmid # 53187; http://n2t.net/addgene:53187; RRID:Addgene_53187, Addgene plasmid # 53188; http://n2t.net/addgene:53188; RRID:Addgene_53188). Then each intact gRNA expression cassette was amplified by PCR and both were inserted into an AAV expression vector that contained a EGFP-KASH reporter (pOTTC1553, Addgene #131682), resulting in plasmid pOTTC1642 (Addgene #195018). All cloning reactions used PCR amplification and ligation-independent cloning (In-Fusion, Takara Inc.). Reactions involving non-AAV plasmids were transformed into Stellar competent cells (Takara Inc.) and those involving AAV plasmids were transformed into NEB Stable competent cells (New England Biolabs).

The pOTTC1642 viral vector was packaged by triple transfection along with pHelper and rep/cap plasmids (serotype 1) and purified by affinity chromatography on GE AVB columns ^60^. Purified vectors were titered by droplet digital PCR with an assay that recognizes the EF1a reporter. Titered vectors were tested for GFP expression by transduction into rat primary cortical neurons prior to *in vivo* use.

#### Immunohistochemistry and *in situ* hybridization

Expression of GRAB-ACh3.0 or eNpHR3.0-eYFP was identified by histological staining for GFP. Briefly, brains slices (40 µm) from perfused (4% paraformaldehyde) brains were washed with 1X PBS, blocked with 1X PBS buffer containing 0.01% Triton X, 10% Normal Donkey Serum and incubated with a primary anti-body for GFP made in chicken (Aves lab, cat#: GFP-1020). Slices were washed and incubated with a secondary anti-chicken antibody made in donkey conjugated to Alexa Fluor 488 (Jackson immuno, cat#: 703-545-155), stained with DAPI and mounted on slides. All images were acquired on a Fluoview 1000 BX51WI confocal microscope (Olympus).

*In situ* hybridization was performed on fresh frozen 16 µm slices according to the ACDBio specifications as described previously ^22^. EGFP-KASH maintained its fluorescence through the protocol. The probes used were as follows: *Chat* (Mm-Chat-C1 (nucleotide target region 1090-1952), GenBank #: NM_009891.2); *Tacr1* (Mm-Tacr1-C3 (nucleotide target region 845-1775), GenBank #: NM_009313.5). All images were taken within a week of the protocol and image intensities were maintained across slices and slides. Quantification was done in the NAc core, where EGFP-KASH expression was found. Due to the cre-independent expression of EGFP-KASH and expression of GFP in ChAT cells, we were unable to specifically identify cells with or without expression and, therefore, all *Chat* positive cells in the expression region were analyzed. RNA was quantified using ImageJ by selecting 10 individual pixel values for every image and averaging to find the intensity values of a single RNA puncta. Intensity values of *Tacr1* in ChAT positive cells were assessed by selecting perinuclear RNA, quantifying intensity, and dividing the total intensity by the intensity of a single puncta value to calculate total RNA puncta. A total of 4 slices from each animal was taken and averaged. Slice averages are represented as individual points.

### QUANTIFICATION AND STATISTICAL ANALYSIS

Detailed statistics can be found in **Table S1**. Data was analyzed and statistics were calculated using Graphpad Prism. Tests were considered significant if *p* < 0.05. For two-sample data, two-sided, un-paired or paired *t*-tests were used. For more than 2 variables over repeated time points one-way or two-way repeated measure (RM) ANOVAs were used followed by post-hoc tests using Sidak’s multiple comparison test. In most cases with an interaction, main effects were reported. For electrophysiology experiments, all are represented as N/n (Mice/cells). Animal exclusions for behavioral experiments were made only when viral expression was absent unilaterally or bilaterally following histological staining. All data is represented as Mean ± standard error. Cartoons were made using BioRender during the subscription period.

## References

1. Zaehle, T., Bauch, E.M., Hinrichs, H., Schmitt, F.C., Heinze, H.-J., Voges, J., Heinze, H.-J., and Bunzeck, N. (2013). Nucleus Accumbens Activity Dissociates Different Forms of Salience: Evidence from Human Intracranial Recordings. The Journal of Neuroscience 33, 8764 LP – 8771. 10.1523/JNEUROSCI.5276-12.2013.

2. Schoenbaum, G., and Setlow, B. (2003). Lesions of Nucleus Accumbens Disrupt Learning about Aversive Outcomes. Journal of Neuroscience 23, 9833–9841.

3. Dutta, S., Beaver, J., Halcomb, C.J., and Jasnow, A.M. (2021). Dissociable roles of the nucleus accumbens core and shell subregions in the expression and extinction of conditioned fear. Neurobiol Stress 15, 100365. 10.1016/j.ynstr.2021.100365.

4. Wendler, E., Gaspar, J.C.C., Ferreira, T.L., Barbiero, J.K., Andreatini, R., Vital, M.A.B.F., Blaha, C.D., Winn, P., and Da, C. (2014). Neurobiology of Learning and Memory The roles of the nucleus accumbens core, dorsomedial striatum, and dorsolateral striatum in learningL: Performance and extinction of Pavlovian fear-conditioned responses and instrumental avoidance responses. Neurobiol Learn Mem 109, 27–36. 10.1016/j.nlm.2013.11.009.

5. Ray, M.H., Russ, A.N., Walker, R.A., and McDannald, M.A. (2020). The Nucleus Accumbens Core is Necessary to Scale Fear to Degree of Threat. The Journal of Neuroscience 40, 4750 LP – 4760. 10.1523/JNEUROSCI.0299-20.2020.

6. Francis, T.C., and Lobo, M.K. (2017). Emerging Role for Nucleus Accumbens Medium Spiny Neuron Subtypes in Depression. Biol Psychiatry 81, 645–653. 10.1016/j.biopsych.2016.09.007.

7. Lee, H.J., Weitz, A.J., Bernal-Casas, D., Duffy, B.A., Choy, M.K., Kravitz, A. v., Kreitzer, A.C., and Lee, J.H. (2016). Activation of Direct and Indirect Pathway Medium Spiny Neurons Drives Distinct Brain-wide Responses. Neuron 91, 412–424. 10.1016/j.neuron.2016.06.010.

8. Kravitz, A. v, Tye, L.D., and Kreitzer, A.C. (2012). Distinct roles for direct and indirect pathway striatal neurons in reinforcement. Nat Neurosci 15, 816–818. 10.1038/nn.3100; 10.1038/nn.3100.

9. Francis, T.C., Chandra, R., Friend, D.M., Finkel, E., Dayrit, G., Miranda, J., Brooks, J.M., Iniguez, S.D., O’Donnell, P., Kravitz, A., et al. (2015). Nucleus accumbens medium spiny neuron subtypes mediate depression-related outcomes to social defeat stress. Biol Psychiatry 77, 212–222. 10.1016/j.biopsych.2014.07.021.

10. Smith, R.J., Lobo, M.K., Spencer, S., and Kalivas, P.W. (2013). Cocaine-induced adaptations in D1 and D2 accumbens projection neurons (a dichotomy not necessarily synonymous with direct and indirect pathways). Curr Opin Neurobiol 23, 546–552. 10.1016/j.conb.2013.01.026.

11. Lobo, M.K., Covington 3rd, H.E., Chaudhury, D., Friedman, A.K., Sun, H., Damez-Werno, D., Dietz, D.M., Zaman, S., Koo, J.W., Kennedy, P.J., et al. (2010). Cell type-specific loss of BDNF signaling mimics optogenetic control of cocaine reward. Science 330, 385–390. 10.1126/science.1188472.

12. Ferguson, S.M., Phillips, P.E., Roth, B.L., Wess, J., and Neumaier, J.F. (2013). Direct-pathway striatal neurons regulate the retention of decision-making strategies. J Neurosci 33, 11668–11676. 10.1523/JNEUROSCI.4783-12.2013; 10.1523/JNEUROSCI.4783-12.2013.

13. Tai, L.H., Lee, A.M., Benavidez, N., Bonci, A., and Wilbrecht, L. (2012). Transient stimulation of distinct subpopulations of striatal neurons mimics changes in action value. Nat Neurosci 15, 1281–1289. 10.1038/nn.3188; 10.1038/nn.3188.

14. Bock, R., Shin, J.H., Kaplan, A.R., Dobi, A., Markey, E., Kramer, P.F., Gremel, C.M., Christensen, C.H., Adrover, M.F., and Alvarez, V.A. (2013). Strengthening the accumbal indirect pathway promotes resilience to compulsive cocaine use. Nat Neurosci 16, 632– 638. 10.1038/nn.3369; 10.1038/nn.3369.

15. Calipari, E.S., Bagot, R.C., Purushothaman, I., Davidson, T.J., Yorgason, J.T., Pena, C.J., Walker, D.M., Pirpinias, S.T., Guise, K.G., Ramakrishnan, C., et al. (2016). In vivo imaging identifies temporal signature of D1 and D2 medium spiny neurons in cocaine reward. Proc Natl Acad Sci U S A 113, 2726–2731. 10.1073/pnas.1521238113 [doi].

16. Iino, Y., Sawada, T., Yamaguchi, K., Tajiri, M., Ishii, S., Kasai, H., and Yagishita, S. (2020). Dopamine D2 receptors in discrimination learning and spine enlargement. Nature 579, 555–560. 10.1038/s41586-020-2115-1.

17. MacPherson, T., Morita, M., Wang, Y., Sasaoka, T., Sawa, A., and Hikida, T. (2016). Nucleus accumbens dopamine D2-receptor expressing neurons control behavioral flexibility in a place discrimination task in the IntelliCage. Learning and Memory 23, 359–364. 10.1101/lm.042507.116.

18. Macpherson, T., Kim, J.Y., and Hikida, T. (2022). Nucleus Accumbens Core Dopamine D2 Receptor-Expressing Neurons Control Reversal Learning but Not Set-Shifting in Behavioral Flexibility in Male Mice. Front Neurosci 16, 885380. 10.3389/fnins.2022.885380.

19. Soares-Cunha, C., Coimbra, B., Domingues, A.V., Vasconcelos, N., Sousa, N., and Rodrigues, A.J. (2018). Nucleus Accumbens Microcircuit Underlying D2-MSN-Driven Increase in Motivation. eNeuro 5, ENEURO.0386-18.2018. 10.1523/ENEURO.0386-18.2018.

20. Soares-Cunha, C., Coimbra, B., David-Pereira, A., Borges, S., Pinto, L., Costa, P., Sousa, N., and Rodrigues, A.J. (2016). Activation of D2 dopamine receptor-expressing neurons in the nucleus accumbens increases motivation. Nat Commun 7, 11829. 10.1038/ncomms11829 [doi].

21. Owesson-White, C., Belle, A.M., Herr, N.R., Peele, J.L., Gowrishankar, P., Carelli, R.M., and Wightman, R.M. (2016). Cue-Evoked Dopamine Release Rapidly Modulates D2 Neurons in the Nucleus Accumbens During Motivated Behavior. J Neurosci 36, 6011– 6021. 10.1523/JNEUROSCI.0393-16.2016.

22. Francis, T.C., Yano, H., Demarest, T.G., Shen, H., and Bonci, A. (2019). High-Frequency Activation of Nucleus Accumbens D1-MSNs Drives Excitatory Potentiation on D2-MSNs. Neuron 103. 10.1016/j.neuron.2019.05.031.

23. Commons, K.G. (2010). Neuronal pathways linking substance P to drug addiction and stress. Brain Res 1314, 175–182. 10.1016/j.brainres.2009.11.014.

24. Nakamura, Y., Izumi, H., Shimizu, T., Hisaoka-Nakashima, K., Morioka, N., and Nakata, Y. (2013). Volume Transmission of Substance P in Striatum Induced by Intraplantar Formalin Injection Attenuates Nociceptive Responses via Activation of the Neurokinin 1 Receptor. J Pharmacol Sci 121, 257–271. 10.1254/jphs.12218FP.

25. Aosaki, T., Tsubokawa, H., Ishida, A., Watanabe, K., Graybiel, A.M., and Kimura, M. (1994). Responses of tonically active neurons in the primate’s striatum undergo systematic changes during behavioral sensorimotor conditioning. J Neurosci 14, 3969– 3984. 10.1523/JNEUROSCI.14-06-03969.1994.

26. Ravel, S., Legallet, E., and Apicella, P. (2003). Responses of Tonically Active Neurons in the Monkey Striatum Discriminate between Motivationally Opposing Stimuli. The Journal of Neuroscience 23, 8489 LP – 8497. 10.1523/JNEUROSCI.23-24-08489.2003.

27. Schulz, J.M., Pitcher, T.L., Savanthrapadian, S., Wickens, J.R., Oswald, M.J., and Reynolds, J.N.J. (2011). Enhanced high-frequency membrane potential fluctuations control spike output in striatal fast-spiking interneurones in vivo. J Physiol 589, 4365– 4381. 10.1113/jphysiol.2011.212944.

28. Aoki, S., Liu, A.W., Zucca, A., Zucca, S., and Wickens, J.R. (2015). Role of Striatal Cholinergic Interneurons in Set-Shifting in the Rat. The Journal of Neuroscience 35, 9424 LP – 9431. 10.1523/JNEUROSCI.0490-15.2015.

29. Okada, K., Nishizawa, K., Fukabori, R., Kai, N., Shiota, A., Ueda, M., Tsutsui, Y., Sakata, S., Matsushita, N., and Kobayashi, K. (2014). Enhanced flexibility of place discrimination learning by targeting striatal cholinergic interneurons. Nat Commun 5, 3778. 10.1038/ncomms4778.

30. Ragozzino, M.E., Mohler, E.G., Prior, M., Palencia, C.A., and Rozman, S. (2009). Acetylcholine activity in selective striatal regions supports behavioral flexibility. Neurobiol Learn Mem 91, 13–22. 10.1016/j.nlm.2008.09.008.

31. Collins, A.L., Aitken, T.J., Huang, I.-W., Shieh, C., Greenfield, V.Y., Monbouquette, H.G., Ostlund, S.B., and Wassum, K.M. (2019). Nucleus Accumbens Cholinergic Interneurons Oppose Cue-Motivated Behavior. Biol Psychiatry 86, 388–396. 10.1016/j.biopsych.2019.02.014.

32. Sandweiss, A.J., and Vanderah, T.W. (2015). The pharmacology of neurokinin receptors in addiction: prospects for therapy. Subst Abuse Rehabil 6, 93–102. 10.2147/SAR.S70350.

33. Tomaz, C., and Nogueira, P.J.C. (1997). Facilitation of memory by peripheral administration of substance P. Behavioural Brain Research 83, 143–145. 10.1016/S0166-4328(97)86058-5.

34. Ebner, K., and Singewald, N. (2006). The role of substance P in stress and anxiety responses. Amino Acids 31, 251–272. 10.1007/s00726-006-0335-9.

35. Aosaki, T., and Kawaguchi, Y. (1996). Actions of substance P on rat neostriatal neurons in vitro. The Journal of neuroscience 16, 5141–5153.

36. Jing, M., Li, Y., Zeng, J., Huang, P., Skirzewski, M., Kljakic, O., Peng, W., Qian, T., Tan, K., Zou, J., et al. (2020). An optimized acetylcholine sensor for monitoring in vivo cholinergic activity. Nat Methods 17, 1139–1146. 10.1038/s41592-020-0953-2.

37. Lubow, R.E. (1973). Latent inhibition. Psychol Bull 79, 398.

38. Soares-Cunha, C., de Vasconcelos, N.A.P., Coimbra, B., Domingues, A.V., Silva, J.M., Loureiro-Campos, E., Gaspar, R., Sotiropoulos, I., Sousa, N., and Rodrigues, A.J. (2020). Nucleus accumbens medium spiny neurons subtypes signal both reward and aversion. Mol Psychiatry 25, 3241–3255. 10.1038/s41380-019-0484-3.

39. Francis, T.C., Chandra, R., Gaynor, A., Konkalmatt, P., Metzbower, S.R., Evans, B., Engeln, M., Blanpied, T.A., and Lobo, M.K. (2017). Molecular basis of dendritic atrophy and activity in stress susceptibility. Mol Psychiatry 22. 10.1038/mp.2017.178.

40. Pignatelli, M., Tejeda, H.A., Barker, D.J., Bontempi, L., Wu, J., Lopez, A., Palma Ribeiro, S., Lucantonio, F., Parise, E.M., Torres-Berrio, A., et al. (2021). Cooperative synaptic and intrinsic plasticity in a disynaptic limbic circuit drive stress-induced anhedonia and passive coping in mice. Mol Psychiatry 26, 1860–1879. 10.1038/s41380-020-0686-8.

41. He, Z.-X., Xi, K., Liu, K.-J., Yue, M.-H., Wang, Y., Yin, Y.-Y., Liu, L., He, X.-X., Yu, H.-L., Xing, Z.-K., et al. (2023). A Nucleus Accumbens Tac1 Neural Circuit Regulates Avoidance Responses to Aversive Stimuli. Int J Mol Sci 24. 10.3390/ijms24054346.

42. Picciotto, M.R., Higley, M.J., and Mineur, Y.S. (2012). Acetylcholine as a neuromodulator: cholinergic signaling shapes nervous system function and behavior. Neuron 76, 116–129. 10.1016/j.neuron.2012.08.036.

43. Oldenburg, I.A., and Ding, J.B. (2011). Cholinergic modulation of synaptic integration and dendritic excitability in the striatum. Curr Opin Neurobiol 21, 425–432. 10.1016/j.conb.2011.04.004.

44. Britt, J.P., Benaliouad, F., McDevitt, R.A., Stuber, G.D., Wise, R.A., and Bonci, A. (2012). Synaptic and behavioral profile of multiple glutamatergic inputs to the nucleus accumbens. Neuron 76, 790–803. 10.1016/j.neuron.2012.09.040; 10.1016/j.neuron.2012.09.040.

45. Gale, G.D., Anagnostaras, S.G., Godsil, B.P., Mitchell, S., Nozawa, T., Sage, J.R., Wiltgen, B., and Fanselow, M.S. (2004). Role of the Basolateral Amygdala in the Storage of Fear Memories across the Adult Lifetime of Rats. The Journal of Neuroscience 24, 3810 LP – 3815. 10.1523/JNEUROSCI.4100-03.2004.

46. Gallo, E.F., Greenwald, J., Yeisley, J., Teboul, E., Martyniuk, K.M., Villarin, J.M., Li, Y., Javitch, J.A., Balsam, P.D., and Kellendonk, C. (2022). Dopamine D2 receptors modulate the cholinergic pause and inhibitory learning. Mol Psychiatry 27, 1502–1514. 10.1038/s41380-021-01364-y.

47. Zhu, Y., Wienecke, C.F.R., Nachtrab, G., and Chen, X. (2016). A thalamic input to the nucleus accumbens mediates opiate dependence. Nature 530, 219–222. 10.1038/nature16954.

48. Al-Hasani, R., Gowrishankar, R., Schmitz, G.P., Pedersen, C.E., Marcus, D.J., Shirley, S.E., Hobbs, T.E., Elerding, A.J., Renaud, S.J., Jing, M., et al. (2021). Ventral tegmental area GABAergic inhibition of cholinergic interneurons in the ventral nucleus accumbens shell promotes reward reinforcement. Nat Neurosci 24, 1414–1428. 10.1038/s41593-021-00898-2.

49. MacAskill, A.F., Cassel, J.M., and Carter, A.G. (2014). Cocaine exposure reorganizes cell type- and input-specific connectivity in the nucleus accumbens. Nat Neurosci 17, 1198– 1207. 10.1038/nn.3783.

50. Matsumoto, N., Minamimoto, T., Graybiel, A.M., and Kimura, M. (2001). Neurons in the thalamic CM-Pf complex supply striatal neurons with information about behaviorally significant sensory events. J Neurophysiol 85, 960–976. 10.1152/jn.2001.85.2.960.

51. Zhang, Y.-F., Reynolds, J.N.J., and Cragg, S.J. (2018). Pauses in Cholinergic Interneuron Activity Are Driven by Excitatory Input and Delayed Rectification, with Dopamine Modulation. Neuron 98, 918–925.e3. 10.1016/j.neuron.2018.04.027.

52. Guzman, R.G., and Kendrick, K.M. (1993). Effect of substance P on acetylcholine and dopamine release in the rat striatum: a microdialysis study. Brain Res 622, 147–154.

53. Brimblecombe, K.R., and Cragg, S.J. (2015). Substance P Weights Striatal Dopamine Transmission Differently within the Striosome-Matrix Axis. Journal of Neuroscience 35, 9017–9023. 10.1523/JNEUROSCI.0870-15.2015.

54. Cachope, R., Mateo, Y., Mathur, B.N., Irving, J., Wang, H.L., Morales, M., Lovinger, D.M., and Cheer, J.F. (2012). Selective activation of cholinergic interneurons enhances accumbal phasic dopamine release: setting the tone for reward processing. Cell Rep 2, 33–41. 10.1016/j.celrep.2012.05.011 [doi].

55. Threlfell, S., Lalic, T., Platt, N.J., Jennings, K.A., Deisseroth, K., and Cragg, S.J. (2012). Striatal dopamine release is triggered by synchronized activity in cholinergic interneurons. Neuron 75, 58–64. 10.1016/j.neuron.2012.04.038 [doi].

56. Kutlu, M.G., Zachry, J.E., Melugin, P.R., Tat, J., Cajigas, S., Isiktas, A.U., Patel, D.D., Siciliano, C.A., Schoenbaum, G., Sharpe, M.J., et al. (2022). Dopamine signaling in the nucleus accumbens core mediates latent inhibition. Nat Neurosci 25, 1071–1081. 10.1038/s41593-022-01126-1.

57. McConalogue, K., Corvera, C.U., Gamp, P.D., Grady, E.F., and Bunnett, N.W. (1998). Desensitization of the neurokinin-1 receptor (NK1-R) in neurons: effects of substance P on the distribution of NK1-R, Galphaq/11, G-protein receptor kinase-2/3, and beta-arrestin-1/2. Mol Biol Cell 9, 2305–2324. 10.1091/mbc.9.8.2305.

58. Bowden, J.J., Garland, A.M., Baluk, P., Lefevre, P., Grady, E.F., Vigna, S.R., Bunnett, N.W., and McDonald, D.M. (1994). Direct observation of substance P-induced internalization of neurokinin 1 (NK1) receptors at sites of inflammation. Proceedings of the National Academy of Sciences 91, 8964–8968. 10.1073/pnas.91.19.8964.

59. Anagnostaras, S., Wood, S., Shuman, T., Cai, D., LeDuc, A., Zurn, K., Zurn, J.B., Sage, J., and Herrera, G. (2010). Automated Assessment of Pavlovian Conditioned Freezing and Shock Reactivity in Mice Using the Video Freeze System. Frontiers in Behavioral Neuroscience 4.

60. Howard, D.B., and Harvey, B.K. (2017). Assaying the Stability and Inactivation of AAV Serotype 1 Vectors. Hum Gene Ther Methods 28, 39–48. 10.1089/hgtb.2016.180.

61. Pennington, Z.T., Dong, Z., Feng, Y., Vetere, L.M., Page-Harley, L., Shuman, T., and Cai, D.J. (2019). ezTrack: An open-source video analysis pipeline for the investigation of animal behavior. Sci Rep 9, 19979. doi: 10.1038/s41598-019-56408-9

