## Supplemental Figures and Tables for "Nucleus Accumbens Local Circuit for Cue-Dependent Aversive Learning"

### Figures S1-6 for Nucleus Accumbens Local Circuit for Cue-Dependent Aversive Learning

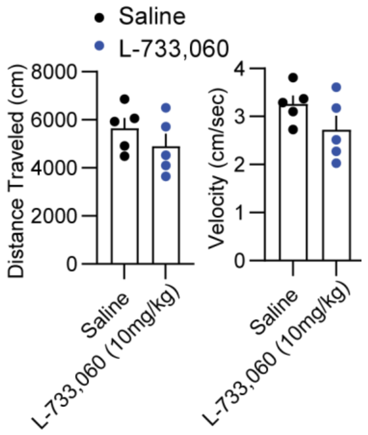


**Figure S1. Neurokinin 1 receptor antagonist L-733,060 does not alter locomotion.** Mice were tested for locomotion in an open field following L-733,060 or saline injection. Open field distance traveled (unpaired two-tailed $t(8) = 1.109$, $p = 0.300$, n = 10 mice) and velocity (unpaired two-tailed $t(8) = 1.577$, $p = 0.154$, n = 10 mice) does not change after an intraperitoneal injection of L-733,060 (10 mg/kg).

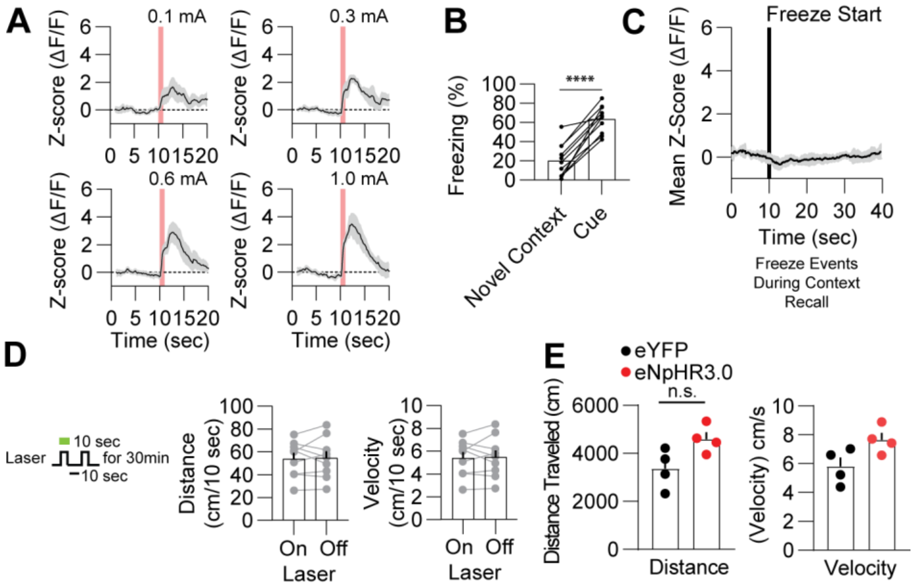


**Figure S2. Dyn-Cre mice freeze to the cue after conditioning and inhibition of D1-MSNs does not affect locomotion. (A)** L-733,060 (i.p.) had no effect on scaling of D1-MSN calcium response to foot shocks of different intensities. **(B)** Dyn-Cre mice show strong freezing compared to the baseline novel context (paired two-tailed $t(10) = 8.911$, $p < 0.0001$, n = 11 mice). **(C)** No change in the D1-MSN calcium signal is observed during freezing events in context recall. **(D)** Dyn-Cre mice underwent 30 min of open field locomotion with halorhodopsin inhibition of D1-MSNs for 10 sec on and 10 sec off. No changes were observed across inhibition and no inhibition periods in distance and velocity measures (paired two-tailed $t(8) = 0.2700$, $p > 0.05$). **(E)** Inhibition of D1-MSNs (8 sec light on 2 sec off for 30 min) in an open field does not alter locomotion measures including distance (unpaired two-tailed $t(6) = 2.396$, $p = 0.054$, n = 8 mice) and velocity (unpaired two-tailed $t(6) = 2.378$, $p = 0.055$, n = 8 mice).

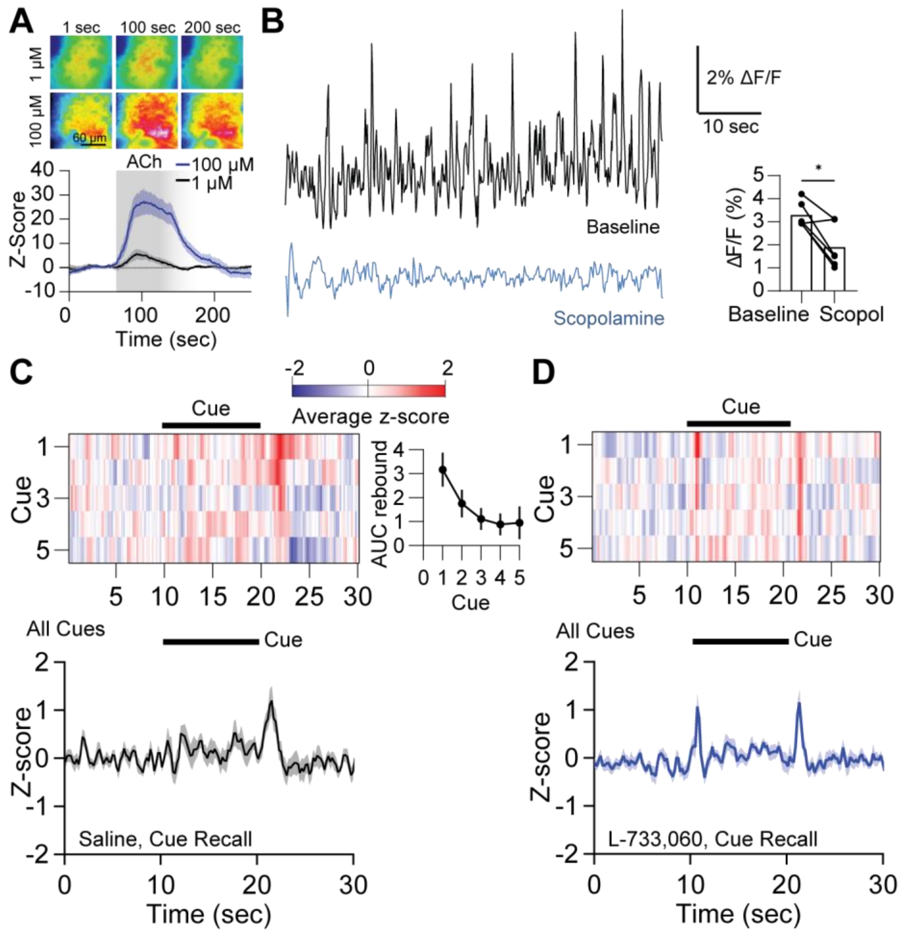


**Figure S3. GRAB-ACh3.0 validation and response to conditioned cues. (A)** Bath application of acetylcholine at 1 µM and 100 µM significantly enhances bulk fluorescence of the GRAB-ACh3.0 sensor in the NAc (Two-way RM ANOVA interaction $F(249, 2490) = p < 0.0001$; concentration main effect $F(1,10) = 15.79$, $p = 0.003$. **(B)** Intraperitoneal injection of scopolamine (1 mg/kg) significantly decreases the averaged *in vivo* acetylcholine signal from GRAB-ACh3.0 in the NAc core ($p < 0.05$, n = 6 mice). A baseline of 2 min was taken immediately after injection of scopolamine while mice were in an open field. The bulk signal (average of the entire signal during this period) was measured. A signal noise floor was calculated and the total signal was calculated as a percentage above the signal floor. **(C-D)** The GRAB-ACh signal was increased following the termination of the cue and this increase diminished over repeated cue exposures (One way RM ANOVA $F(4,28) = 4.011$, $p = 0.0392$, n = 8). However, the ACh signal was also increased during the start of the cue in mice injected with L-733,060 during conditioning.

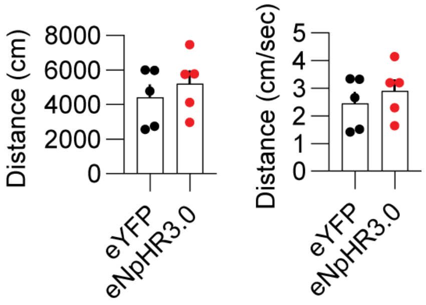


**Figure S4. ChAT inhibition has no effect on locomotion.** No difference in distance traveled (two-tailed unpaired $t(8) = 0.7431$, $p = 0.4787$, n = 5, 5) or velocity (two-tailed unpaired $t(8) = 0.7431$, $p = 0.4787$, n = 5, 5) was observed during halorhodopsin inhibition (8 sec light on 2 sec off for 30 min) of ChAT neurons in the open field.

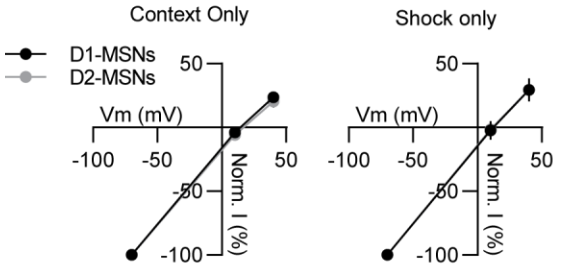


**Figure S5. Electrophysiological changes following conditioning are selective to D2-MSNs.** No differences in I-V rectification are observed between D1-MSNs and D2-MSNs in a context only condition (Two-way RM ANOVA $F(2,8) = 0.122$, $p = 0.887$, n = 6 cells, 3 mice) and a shock only condition (Two-way RM ANOVA $F(2,4) = 0.005$, $p = 0.995$, n = 4 cells, 2 mice).

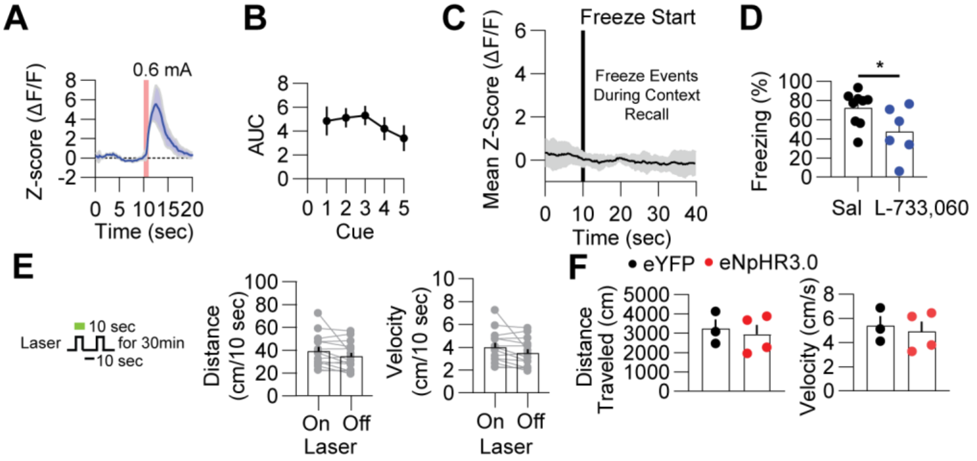


**Figure S6. NAc core D2-MSN-related responses to shock and optogenetic manipulation. (A)** L-733,060 injection had no effect on the size of the shock-induced calcium response in D2-MSNs. **(B)** Area under the curve (AUC) of GCaMP7f fluorescent response to cues does not vary during cue recall (One-way ANOVA $F(4, 20) = 0.6139$, $p = 0.6575$). **(C)** No D2-MSN calcium response to individual freeze events is observed during context recall. **(D)** Freezing to the conditioned cue was significantly decreased in A2a-cre photometry mice treated with L-733,060 (two-tailed unpaired $t(13) = 2.180$, $p = 0.0483$, n = 9, 6). **(E)** A2a-Cre mice underwent 30 min of open field locomotion with halorhodopsin inhibition of D2-MSNs for 10 sec on and 10 sec off. No changes were observed across inhibition and no inhibition periods in distance and velocity measures (two-tailed paired $t(14) = 1.882$, $p > 0.05$, n =15). **(F)** No difference in distance traveled (two-tailed unpaired $t(5) = 0.394$, $p = 0.710$, n = 7) or velocity (two-tailed unpaired $t(5) = 0.394$, $p = 0.709$, n = 7) was observed during inhibition (8 sec light on 2 sec off for 30 min) of D2-MSNs in the open field.

**Table S1: Statistics**

| Figure | Panel | Test | Comparison | Test Statistic | Exact *P*-value |
|---|---|---|---|---|---|
| 1 | B | Two-tailed unpaired *t*-test | Context | *t*(40) = 0.398 | 0.6928 |
| | | Two-tailed unpaired *t*-test | Cue | *t*(40) = 3.801 | 0.0005 |
| | C | Two-tailed unpaired *t*-test | Context | *t*(13) = 0.539 | 0.5882 |
| | | | Cue | *t*(13) = 1.337 | 0.2040 |
| | D | Two-tailed unpaired *t*-test | ChAT+ | *t*(6) = 5.315 | 0.0009 |
| | | Two-tailed unpaired *t*-test | ChAT- | *t*(6) = 0.955 | 0.1883 |
| | E | Two-way RM ANOVA | | Interaction *F*(6,60) = 7.442 | <0.0001 |
| | | | Baseline vs. SP: ChAT-Cre/Cas9$^{Tac1r\text{-}gRNAs}$ | Sidak's mc *t*(60) = 1.669 | 0.1906 |
| | | | ChAT-Cre$^{Tac1r\text{-}gRNAs}$ | Sidak's mc *t*(60) = 7.090 | <0.0001 |
| | | | ChAT-Cre | Sidak's mc *t*(60) = 7.315 | <0.0001 |
| | | | ChAT-Cre/Cas9 | Sidak's mc *t*(60) = 6.518 | <0.0001 |
| | F | Two-tailed unpaired *t*-test | Context | *t*(26) = 0.028 | 0.9780 |
| | | Two-tailed unpaired *t*-test | Cue | *t*(26) = 4.419 | 0.0003 |
| | G | Two-tailed unpaired *t*-test | Context | *t*(13) = 0.442 | 0.6656 |
| | | Two-tailed unpaired *t*-test | Cue | *t*(13) = 4.073 | 0.0013 |
| 2 | C | One-way ANOVA | | Interaction *F*(3,33) = 28.40 | <0.0001 |
| | | | 0.1 mA vs. 1.0 mA | Sidak's mc *t*(33) = 8.727 | <0.0001 |
| | D | One-way RM ANOVA | | *F*(9, 81) = 1.281 | 0.2601 |
| | G | One-way ANOVA | 0.1 mA vs. 1.0 mA | *F*(3, 28) = 18.81 | <0.0001 |
| | H | Pearson r | | r(22) = 0.5525 | 0.0077 |
| | J | Two-tailed unpaired *t*-test | Context | *t*(19) = 0.869 | 0.3959 |
| | | Two-tailed unpaired *t*-test | Cue | *t*(19) = 2.989 | 0.0075 |
| | K | Two-tailed unpaired *t*-test | Context | *t*(17) = 1.162 | 0.2611 |
| | L | Two-way RM ANOVA | Time course | No Interaction *F*(6, 102) = 1.736 | 0.1202 |
| | | Two-way RM ANOVA | Group data: Laser On vs. Off | No Interaction *F*(1, 17) = 0.574 | 0.4592 |
| 3 | C | Mixed-effects analysis | Pause | No Interaction *F*(3,62) = 0.031 | 0.9923 |

| | | | | | |
|---|---|---|---|---|---|
| | D | Mixed-effects analysis | Rebound | Interaction $F(3,62) = 2.795$ | 0.0476 |
| | | | Shock trial 1 vs. Shock trial 10 | Main effect $F(1, 62) = 24.56$ | <0.0001 |
| | F | Two-way RM ANOVA | Pause | No Interaction $F(1,20) = 1.225$ | 0.2830 |
| | G | Two-way RM ANOVA | Rebound | Interaction $F(1,20) = 4.389$ | 0.0491 |
| | | | Shock trial 1, Saline vs. L-733,060 | Sidak's mc $t(20) = 3.284$ | 0.0221 |
| | J | Mixed-effects analysis | | Interaction $F(1,36) = 6.221$ | 0.0174 |
| | K | Pearson r | | $r(12) = 0.7228$ | 0.0079 |
| 4 | B | Two-tailed unpaired *t*-test | Context | $t(10) = 1.798$ | 0.1025 |
| | | Two-tailed unpaired *t*-test | Cue | $t(10) = 0.5110$ | 0.6205 |
| | C | Two-tailed unpaired *t*-test | Context | $t(12) = 1.583$ | 0.1394 |
| | | Two-tailed unpaired *t*-test | Cue | $t(12) = 4.395$ | 0.0009 |
| | D | Two-tailed unpaired *t*-test | Context | $t(17) = 0.135$ | 0.8941 |
| | | Two-tailed unpaired *t*-test | Cue | $t(17) = 3.253$ | 0.0047 |
| | E | Two-tailed unpaired *t*-test | Context | $t(11) = 1.153$ | 0.2734 |
| | | Two-tailed unpaired *t*-test | Cue | $t(11) = 1.220$ | 0.2479 |
| 5 | A | Two-way RM ANOVA | IV Curve | Interaction $F(2, 34) = 8.820$ | 0.0008 |
| | | | IV: +40 mV | Sidak's mc $t(51) = 4.691$ | <0.0001 |
| | | One-way ANOVA | Rectification Index | $F(2, 24) = 7.194$ | 0.0036 |
| | B | Two-tailed unpaired *t*-test | 20-25 min | $t(9) = 2.444$ | 0.0371 |
| | C | One-way ANOVA | sEPSC amp | $F(2, 27) = 5.177$ | 0.0125 |
| | | | sEPSC amp, Cue vs. Cue + Shock | Sidak's mc $t(22) = 3.171$ | 0.0075 |
| | | One-way ANOVA | sEPSC freq | $F(2, 27) = 0.6541$ | 0.5280 |
| | D | Two-way RM ANOVA | IV Curve | $F(4, 36) = 0.5051$ | 0.7322 |
| | | One-way ANOVA | Rectification Index | $F(2,19) = 1.491$ | 0.2503 |
| 6 | C | Mixed-effects model | Shock Intensity | $F(3, 38) = 1.656$ | 0.1928 |
| | D | One-way RM ANOVA | Shock Event | $F(9, 81) = 1.300$ | 0.2499 |
| | I | One-way ANOVA | | $F(2, 24) = 7.592$ | 0.0028 |
| | | Dunnet mc | D1-MSN vs. D2-MSN | $q(24) = 3.643$ | 0.0025 |

| | | | | | |
|---|---|---|---|---|---|
| | | Dunnet mc | D2-MSN vs. D2-MSN L-733,060 | *q*(24) = 3.125 | 0.0086 |
| | J | Pearson r | | *r*(16) = 0.3768 | 0.0201 |
| | L | Two-tailed unpaired *t*-test | Context | *t*(14) = 2.527 | 0.0242 |
| | M | Two-way RM ANOVA | Over Cues | Interaction *F*(6, 90) = 3.905 | 0.0016 |
| | | Two-way RM ANOVA | Group data: Laser On vs. Off | Interaction *F*(1, 14) = 13.57 | 0.0025 |
| | | Uncorrected Fisher's LSD | Off: eYFP vs. eNpHR3.0 | *t*(28) = 1.642 | 0.1117 |
| | | Uncorrected Fisher's LSD | On: eYFP vs. eNpHR3.0 | *t*(28) = 3.784 | 0.0007 |
| | N | Two-tailed unpaired *t*-test | Context | *t*(16) = 0.3347 | 0.3347 |
| | O | Two-way RM ANOVA | Over cues | Interaction *F*(6, 96) = 3.153 | 0.0073 |
| | | Two-way RM ANOVA | Group data: Laser On vs. Off | Interaction *F*(1, 13) = 33.63 | <0.0001 |
| | | | On: eYFP vs. eNpHR3.0 | Sidak's mc *t*(26) = 2.526 | 0.0356 |
